# Generation of a molecular interactome of the glioblastoma perivascular niche reveals Integrin Binding Sialoprotein as a key mediator of tumor cell migration

**DOI:** 10.1101/2021.10.01.462643

**Authors:** Yasmin Ghochani, Alireza Sohrabi, Sree Deepthi Muthukrishnan, Riki Kawaguchi, Michael C. Condro, Soniya Bastola, Fuying Gao, Yue Qin, Jack Mottahedeh, M. Luisa Iruela Arispe, Nagesh Rao, Dan R. Laks, Linda M. Liau, Gary W. Mathern, Steven A. Goldman, S. Thomas Carmichael, Ichiro Nakano, Giovanni Coppola, Stephanie Seidlits, Harley I. Kornblum

**Affiliations:** Department of Psychiatry and the Semel Institute for Neuroscience and Behavior, David Geffen School of Medicine at UCLA; Department of Bioengineering, UCLA; Department of Neurology, David Geffen School of Medicine at UCLA; Department of Cell and Developmental Biology, Northwestern University; Department of Pathology and Laboratory Medicine, David Geffen School of Medicine at UCLA; Voyager Therapeutics, Cambridge, MA 02139; Department of Neurosurgery, David Geffen School of Medicine at UCLA; Department of Neurology and the Center for Translational Neuromedicine, University of Rochester; Research and Development Center for Precision Medicine, Tsukuba University, Japan; Departments of Pediatrics and Pharmacology, David Geffen School of Medicine, at UCLA

## Abstract

Glioblastoma (GBM) is characterized by extensive microvascular hyperproliferation. In addition to supplying blood to the tumor, GBM vessels also provide trophic support to glioma cells and serve as conduits for migration into the surrounding brain promoting recurrence. Here, we enriched CD31-expressing glioma vascular cells (GVC) and A2B5-expressing glioma tumor cells (GTC) from primary GBM and utilized RNA sequencing to create a comprehensive interaction map of the secreted and extracellular factors elaborated by GVC that can interact with receptors and membrane molecules on GTC. To validate our findings, we utilized functional assays, including a novel hydrogel-based migration assay and in vivo mouse models to demonstrate that one identified factor, the little-studied integrin binding sialoprotein (IBSP) enhances tumor growth and promotes the migration of GTC along the vasculature. This perivascular niche interactome will serve a resource to the research community in defining the potential functions of the GBM vasculature.

## Introduction

Glioblastoma (GBM) is the most common primary brain tumor and is virtually always fatal, universally recurring following the standard of care therapies of surgery, radiation and chemotherapy with Temozolomide (Stupp et al., 2015). One of the primary hallmarks of GBM is extensive microvascular proliferation, characterized by dysfunctional and atypical vessels with variable diameter and permeability and heterogeneous distribution (Brat et al., 2004) (Das and Marsden, 2013).

The GBM microvasculature largely consists of endothelial cells, pericytes, and vascular-associated immune cells that play a critical role in the maintenance of blood flow to the tumor cells (Charles and Holland, 2010). There is increasing evidence that vasculature can play other important roles in GBM. For example, vascular cells are known to provide a means by which GBM cells migrate out of the tumor, which then serve as the seeds of recurrence following therapy (Farin et al., 2006; Griveau et al., 2018). In experimental models, vascular cells have been demonstrated to provide trophic support to tumor cells, outside of their role in supplying blood, including allowing them to survive therapeutic insults such as radiation (Garcia-Barros et al., 2003; Calabrese et al., 2007; Charles et al., 2010; Yan et al., 2014; Brooks and Parrinello, 2017).

While a great deal is known about how GBM cells induce the ingrowth and elaboration of the GBM vasculature, few studies have highlighted the factors that the microvasculature produces factors that may act on the tumor cells, and how tumor cells may respond to these factors (Bao et al., 2006; Brat and Van Meir, 2001; Gilbertson and Rich, 2007). In order to investigate the potential tumor-promoting roles of the microvasculature, early studies have tried to identify a core set of dysregulated genes associated with aberrant GBM vessels as compared to normal brain vessels (Charalambous et al., 2005, 2007; Pen et al., 2007; Dieterich et al., 2012). Although these studies provided a great deal of information, due to technical limitations, they did not yield a comprehensive map of the factors elaborated by GBM vasculature.

In this study, we have overcome some of these technical limitations by performing RNA sequencing of highly enriched, primary GBM vascular and tumor cells. We have created an extensive map of interactions among potentially secreted and extracellular factors elaborated by the GBM vascular cells (GVC) and receptors and other membrane-bound molecules on GBM tumor cells (GTC), which represents a new resource for the research community. To validate our findings, we demonstrate that one of our identified factors, the little-studied Integrin Binding Sialoprotein (IBSP), provides trophic support and promotes the migration of GTC along the vasculature.

## Results

### Transcriptomic profiling of cells expressing CD31 or A2B5 from non-transformed cortices and primary GBM uncovers tumor-specific dysregulated genes

To delineate the molecular interactions between the perivascular niche (PVN) and tumor cells, we developed a strategy involving the sequential enrichment of CD31+ (PECAM1) vascular cells (GVC) and A2B5+ tumor cells (GTC) from primary GBM samples. We utilized these markers because virtually all endothelial cells are CD31+, and the vast majority of tumor-initiating cells are contained within the A2B5+ fraction (Auvergne et al., 2013; Tchoghandjian et al., 2010). This A2B5+ fraction harbors the major chromosomal alterations of the parent tumor (**Figure S1A**). These markers were also used to enrich for their non-neoplastic counterparts isolated from pathologically normal regions of tissue resected for chronic epilepsy - normal brain glial progenitor cells (BGPC) and normal brain vascular cells (BVC), allowing direct comparisons (**Figure 1A**). Information on specific patient diagnoses and sample characteristics are described in **Tables S1A** (non-transformed) and **Table S1B** (GBM).

**Figure 1:**
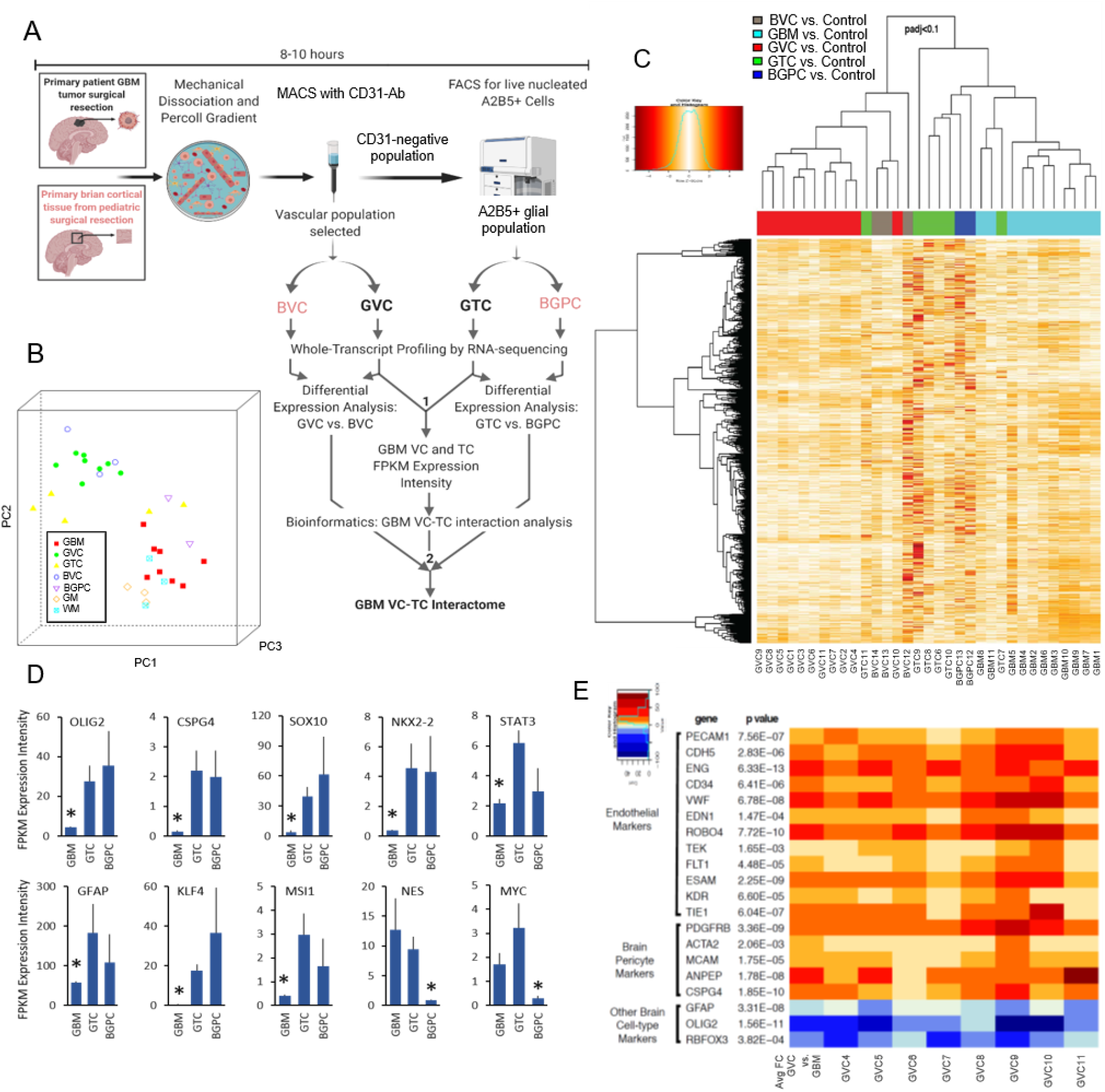
Transcriptomic characterization of GBM tumor and vascular cells. (A) Experimental approach for isolation and characterization of tumor and non-neoplastic cells from GBM patient and brain cortices (B) PCA of un-dissociated primary GBM samples (red), and GTC (yellow) and GVC (green) derived from them, along with non-transformed BGPC (purple), BVC (blue), GM (orange), and WM (aqua). (C) Heatmap shows gene expression profile and clustering of all 33 samples (D) Marker expression analysis GTC vs GBM and GTC vs. BGPC. * FDR<0.05 (E) Vascular cell marker enrichment analysis. Vascular markers upregulated in MACS-enriched GVC as compared to the whole un-dissociated parent tumor (GBM) obtained through paired differential expression analysis (p<0.005). See also Figure S1

Principal-component analysis (PCA) of gene expression profiles of all samples showed close clustering of the GVC, and BVC fraction, and variability in GTC and BGPC transcriptomes (**Figure 1B**). Differential gene expression analysis comparing each sample vs combined expression of non-neoplastic gray and white matter revealed similarity in expression profiles within each cellular fraction, while distinguishing each individual fraction (**Figure 1C**). Specifically, examination of differential gene expression analysis of the GTC vs. BGPC identified 1471 transcripts that were significantly enriched and 1092 that were significantly depleted in GTC (**File S1A**). We utilized Ingenuity Pathway Analysis (IPA) to identify significantly enriched diseases or functions associated with the differentially expressed genes in GTC vs. BGPC fractions. Significant upregulation of tumor pathways promoting proliferation, survival, CNS neoplasia and gliomagenesis, and downregulation of normal brain functions such as learning and cognition supported the neoplastic nature of the isolated GTC, and the inclusion of the tumor propagating cells within the A2B5+ fraction **(Figure S1B, File S1B**).

We further evaluated the expression of known astroglial- and oligodendroglial-progenitor markers, neural stem/progenitor cell markers, and other regulators of cancer stem cells using differential expression and direct expression intensity analyses (FPKM values, raw data in **Figure S1C-D**, respectively). GTC were enriched for the oligodendroglial progenitor markers Olig2, CSPG4, SOX10, and NKX2-2 as compared to the unsorted parent tumors and, as expected, these markers were also enriched in BGPC (**Figure 1D**). Interestingly, GTC also exhibited significant upregulation of STAT3 and GFAP, an astrocytic marker, and progenitor cell markers KLF4 and MSI1 as compared to unsorted GBM. BGPC, however, did not significantly exhibit these markers as compared to GBM samples or their A2B5+ fractions. Lastly, both the parent GBM and their GTC fractions had significantly elevated expression of the neural stem cell marker NES, and cancer stem cell regulator MYC as compared to the BGPC, consistent with the hypothesis that the GTC exhibit multipotent tumor stem/progenitor-like character. Taken together, these findings strongly support the notion that the A2B5+ fraction is highly enriched for tumor cells, and that the cells contained within this fraction may have stem-like qualities.

### Molecular characterization and identification of GVC-specific factors

To validate the identity of our CD31-enriched GBM vascular cell (GVC) fraction, we examined endothelial and pericyte markers both of which were significantly upregulated in GVC samples vs. the parent tumors supporting a mixed endothelial/pericyte origin (**Figure 1E**). As expected, GVC had significantly downregulated expression of the astrocyte, oligodendrocyte, and neuronal markers GFAP, OLIG2, and RBFOX3 (NeuN), respectively. The presence of pericyte-like markers within this fraction could indicate that pericytes remained tightly bound to endothelial cells during the enrichment process or that some genes generally associated with pericytes are expressed by CD31+ GVC.

We identified 445 genes enriched in GVC compared to BVC (p< 0.005) and capturing known regulators of the GBM-perivascular interaction such as ANGPT2 (Scholz et al., 2016; Stratmann et al., 1998), VEGF-A induced endothelial genes ESM1, NOX4, PXDN (Dieterich et al., 2012), and TGFβR1 (Krishnan et al., 2015), along with many novel putative targets (**Figure 2A, File S2a**). In addition, examination of the significantly activated upstream regulators of GVC revealed not only the known vascular regulators in GBM, such as TGFβ, VEGF, HIF1α, NOS2, various RTKs, and SPP1 and Endothelin-1 (EDN1, p-value=0.01) (Dieterich et al., 2012; Jeon et al., 2014; Musumeci et al., 2015), but also novel putative regulators such as IL1A/B (IL1Ap-value=1.6×10^−5^; IL1B:, p-value=7.8×10^−7^) (**Figure 2A, File S2b**). IPA analysis of GVC vs. BVC transcriptomes revealed significant enrichment of GO terms related to glioma invasiveness and proliferation, and negative association with tumor necrosis and cell death (**Figure 2B, File S2B**,**C**). Thus, our unbiased transcriptomic analysis of the freshly isolated GVC provides the opportunity for investigation of multiple dysregulated genes and pathways within the GBM PVN.

**Figure 2:**
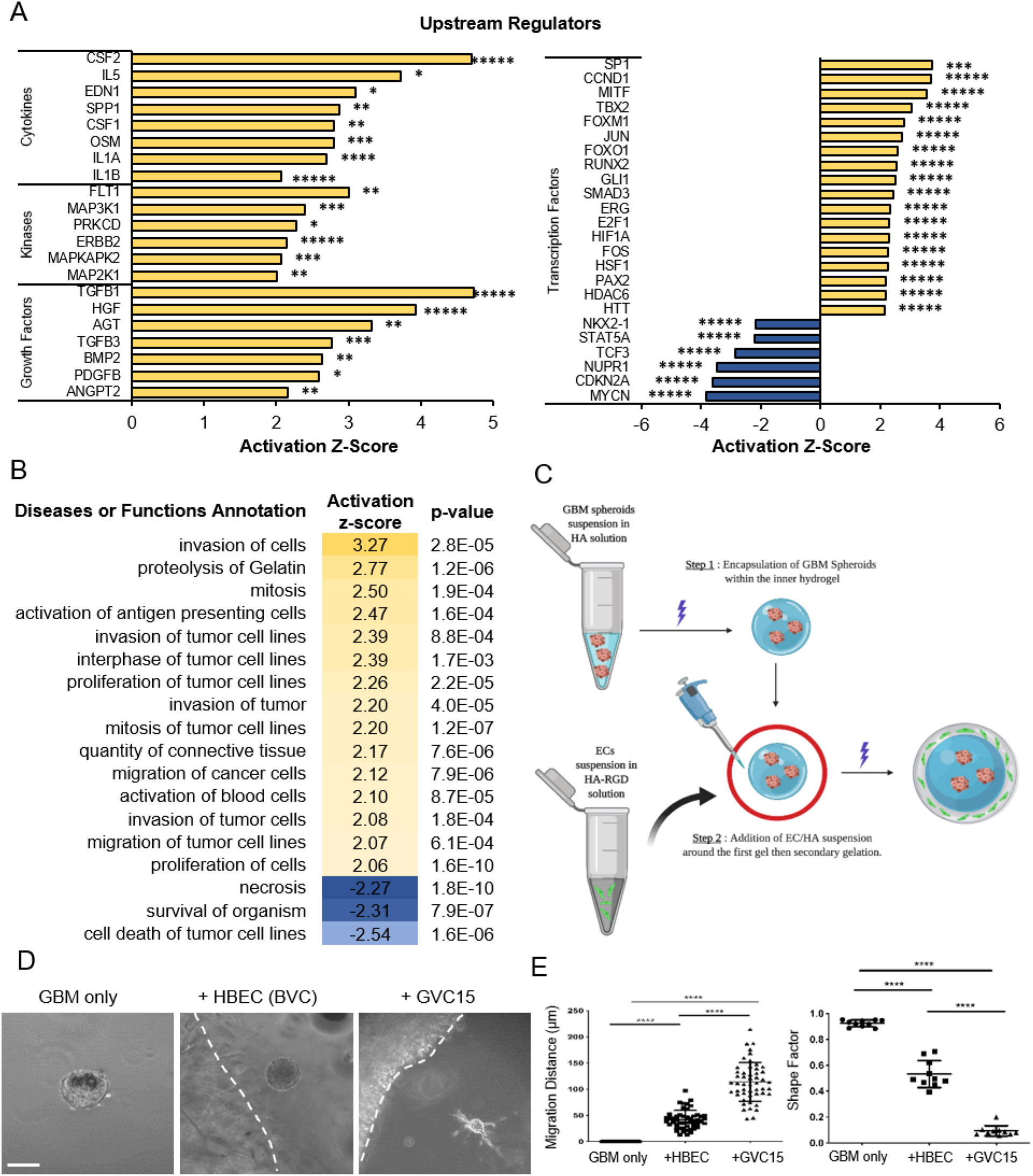
Enrichment of migration-related function and pathways in GVC transcriptome. (A, B) IPA analysis shows enrichment of various growth factors, signaling pathways and disease or functions in GVC (C) Schematic shows the strategy for assessing migration in hydrogels encapsulated with GBM and GVC. (D-F) Images show migration of GBM spheroids alone or with HBEC (BVC) or GVC encapsulated in hydrogels. Graphs show quantitation of migration distance and shape factor. N=10 GBM spheroids, Mann Whitney test, * * p<0.0001. Scale bar,100um. See also Figure S2

### Generation of the GTC-GVC molecular interactome reaffirms a prominent role for perivascular niche-regulation of GBM migration

Because of the high degree of enrichment of GO terms associated with invasiveness and migration, we sought to determine whether GVC-secreted factors could directly influence invasive capacity of GBM cells. To accomplish this, we developed a novel 3D hydrogel-based system in which we separately encapsulated GBM gliomaspheres and short-term primary cultures of GVC or BVC, placing them in close proximity to each other (**Figure 2C**). The vascular characteristics of GVC were confirmed by CD31 immunostaining, a Di-Ac-LDL uptake assay and gene expression profiling (**Figure S2A-C**). Results indicated a probable mixed endothelial/pericyte identity. Gliomaspheres cultured alone or in presence of BVC demonstrated different migration potential (measured as the distance of cell movement away from the sphere edge), with no migration (0±0 µm) or minimal migration (39.9±21.8 µm) (p<0.0001) and shape factor analysis (where 1 indicates no migratory potential and 0 indicates high migratory potential) revealed non-migratory (0.9±0.02) or mildly migratory (0.5±0.1) (p<0.0001) (**Figure 2D, E**). However, gliomaspheres co-encapsulated with GVC showed significantly increased average migration distance of 116.2±30.4 µm (p<0.0001) and aggressive migratory potential with shape factor average of 0.09±0.04 (p<0.0001) (**Figure 2D, E**). These findings support the hypothesis that a factor or factors elaborated by GVC promote GBM migration.

Given that our functional 3D *in vitro* assays demonstrated increased GBM migratory potential in the presence of GVC-secreted factors, and our transcriptomic analysis showed enrichment of pro-migration/invasion-related pathways, we sought to identify specific angiocrine signaling cues elaborated by GBM vasculature that can influence tumor cell biology. A putative GTC-GVC molecular interactome was developed by generating unique transcriptional profiles to identify specific cues in GVC that can mediate the pro-migratory effect on GTC. We began our analysis by scrutinizing the 113 GVC extracellular factors and the 331 GTC plasma membrane (PM) proteins, exclusively, which were differentially regulated as compared to BVC and BGPC. Differential expression of angiocrines in GVC vs BVC are shown in **File S3A**, and differential expression of GTC vs. BGPC plasma membrane (PM) proteins are shown in **File S3B**. Utilizing a combination of manual and IPA software curation focused on previously reported direct and indirect protein-protein interactions, we identified 24 interacting groups encompassing hormones, extracellular matrix components such as laminins, collagens, and matrix metalloproteinases, and members of the small integrin-binding ligand N-linked glycoprotein (SIBLING) family of proteins (**Figure 3A**). In the case of the SIBLING proteins, the interaction partners are mostly those previously reported for the osteopontin (OPN)-encoding gene, SPP1 (Lamour et al., 2015). However, due to conservation of various functional motifs, such as post-translational modification motifs, acidic amino acids, and the RGD (Arg-Gly-Asp) motif, as well as the much more prominent and significant upregulation of another member of this family, IBSP as compared to SPP1, we extended the interaction unit to include all upregulated members of the SIBLING family (Bellahcène et al., 2008). As expected, IPA of the diseases and functions associated with the genes within this interactome revealed cellular movement and migration as the most significantly upregulated, followed by angiogenesis, and cellular proliferation (**Figure 3B**).

**Figure 3:**
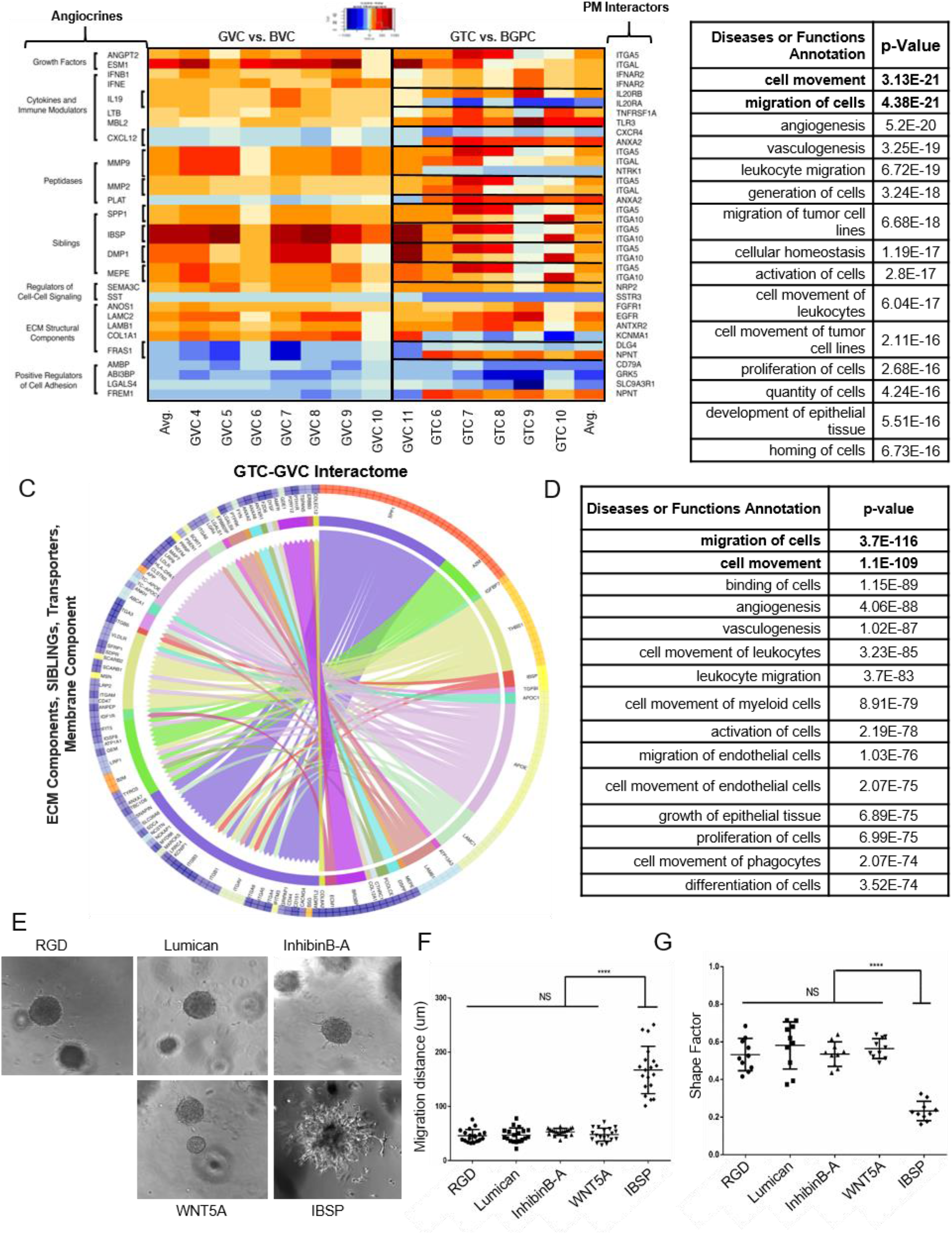
Generation of a perivascular niche-tumor interactome. (A) Heatmap shows differentially regulated genes associated with putative GVC angiocrine-GTC plasma membrane compared to non-transformed BVC and BGPC. (B) IPA diseases and functions significantly associated with the genes in the interactome (C) Representative circos plot of the interactome comprising of ECM Components, SIBLINGs, Transporters, Membrane Components. (D) IPA diseases or functions significantly associated with all angiocrines and their putative GTC-PM interaction partners. (E-G) Representative images of GBM spheroids encapsulated in hydrogels treated with the indicated factors. Scale bar,100um. Graphs show quantitation of shape factor and migration distance. N=10 GBM spheroids, Mann Whitney test, * * p<0.0001. See also Figure S3

The interactome described above, assumed that both the extracellular factors/ligands and the plasma membrane proteins/receptors must necessarily be dysregulated for their interaction to be of consequence in promoting tumor migration. In order to obtain a more comprehensive view of all of the GVC angiocrines and their putative interaction with all the GTC membrane proteins, we utilized FPKM expression intensity values and considered those with FPKM>1 for at least 1/3 of the samples to account for the heterogeneity that exists among GBM. We identified 552 GVC extracellular factors and 1254 GTC plasma membrane genes. GVC extracellular factors FPKM (a), GTC PM proteins FPKM (b), and our comprehensive PVN interactome by FPKM expression units (c) are shown in **File S4**, where FPKM>4 are highlighted in red (collagen and complement factor interactions are not shown). We eliminated factors that were expressed by both GVC and GTC with an FPKM difference of <4, thus narrowing the interactome to putative GTC dependence on GVC, and not GTC autocrine signaling. We identified 135 angiocrines that had at least one, but often multiple, putative GTC-interacting partners. Grouped into different functional categories, the Circos plots demonstrate the angiocrines on the right hemi-circle and their color-coded GTC interacting partners on the left for easy identification of known and novel GTC-GVC tumor-angiocrine interactions, with the outer circles demonstrating expression levels of various factors (**Figure S3A**). A representative plot depicting the ECM components, SIBLINGS, Transporters, Membrane Components is shown in **Figure 3C**.

IPA annotation of functions associated with this comprehensive interactome (**Figure 3D, File S4**) confirmed that the most significantly enriched GO_Term is cellular migration and movement, again indicating that promotion of tumor invasion is a likely function of the interacting proteins. Furthermore, there was a modest enrichment of terms associated with tumor proliferation, suggesting a potential role for the vascular angiocrines in this process. Within this comprehensive interactome, we found many previously reported PVN angiocrine-receptor interactions validating our experimental approach (**Figure S3B**). In addition, we identified multiple novel putative interactions such as Inhibin angiocrines of the TGFβ superfamily, ECM small leucine-rich proteoglycan (SLRP) family of proteins like Lumican and Biglycan, and Nidogen angiocrine basement membrane components, along with their putative binding partners demonstrating the complexity of the signaling emanating from the PVN (**Figure 3C**).

### IBSP regulates tumor cell migration, proliferation and induces a mesenchymal signature

Since tumor invasion and migration-related signaling pathways were highly enriched within the PVN interactome, we tested a few select candidate angiocrines: Lumican, InhibinB-A, WNT5A, and IBSP in promoting migration of GTC using gliomaspheres derived from a patient line (HK_217) in hyaluronic acid (HA)-based hydrogel system. Since the GBM microenvironment is rich in RGD-containing factors, which are necessary for migration, and GTC express high levels of integrin receptors, we used an RGD peptide-conjugated HA culture scaffold as our positive control, and all factors were studied on the background of RGD-conjugated hydrogels, using a cysteine-only hydrogel with no integrin-binding sites as a negative control (**Figure 3E**). Of the angiocrines assessed, IBSP (Integrin binding Sialoprotein, BSPII) peptide was the only factor that promoted significant migration above the baseline distance promoted by the RGD control (**Figure 3E-F**).

Within the PVN interactome, multiple SIBLING family members were highly upregulated in GVC as compared to BVC. However, IBSP was the most upregulated member of this family (logFC of 13, p=0.0001) and almost exclusively expressed by GVC (FPKM=50.1 vs. GTC FPKM=2.4) (**Figure S4A-C**). Immunostaining of multiple primary GBM tissues validated the vascular expression of IBSP protein (**Figure 4A, Figure S4D**). Vascular expression of IBSP transcript was also confirmed by fluorescent *in situ* hybridization, although the precise cell types that expressed IBSP mRNA could not be easily determined with some apparent co-expression in PECAM-1 mRNA-expressing endothelial cells and PDGFRβ mRNA-expressing presumptive pericytes (**Figure 4B**).

**Figure 4:**
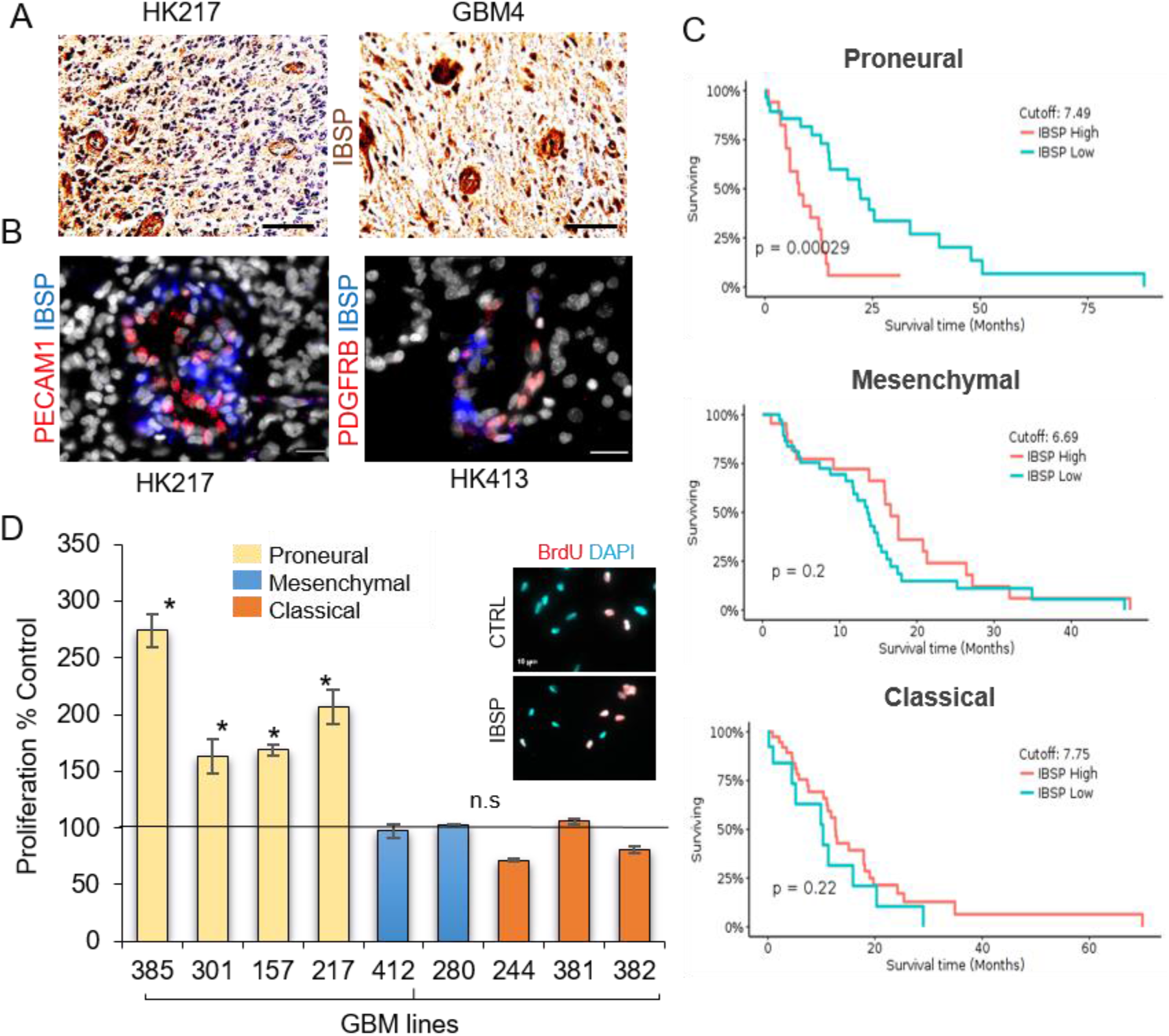
IBSP expression correlates with poor survival in Proneural GBM. (A) Immunohistochemistry for IBSP (brown) in 3 GBM patient tumor tissue. Nuclei counterstained with hematoxylin (blue). Scale bars, 100um (B) Representative images of in situ for IBSP with PECAM1 (Endothelial) and PDGFRB (pericyte) markers in GBM tissue. Scale bars, 20um (C) Kaplan-Meier plots of patient survival correlation with IBSP gene expression in TCGA samples from each molecular subtype. (D) Graph shows proliferation of GBM cells from the 3 TCGA molecular subtypes treated with IBSP. **\*** p<0.05 one-tailed paired t-test. Images show representative BrdU staining in CTRL and IBSP treated cells. Scale bars, 10um. See also Figure S4.

To determine whether IBSP expression was associated with patient outcome, we utilized the open-access Gliovis tool to assess GBM patient survival correlation with IBSP expression within different tumor subtypes (Bowman et al., 2017). We found that elevated IBSP correlated with worse survival only in the Proneural subtypes (p=0.0003) (**Figure 4C**). Because of the enrichment of terms associated with proliferation of tumor cells, we added IBSP to gliomasphere cultures from all 3 subtypes, and found elevated cell numbers only in the gliomaspheres that had previously been characterized as “Proneural” (**Figure 4D**), accompanied by an increase in the BrdU incorporation (**Figure 4D, inset**). These findings indicate that IBSP could play an important role in regulating proliferation of Proneural GBM.

To determine the molecular basis of IBSP-induced effects, we treated 2 Proneural GBM cultures with IBSP and profiled gene expression by cDNA microarray. Differential gene expression analysis revealed upregulation of genes associated with Mesenchymal phenotype such as CD44 (p< 2.26E-09) and MMP1 (p<3.03E-11) in IBSP-treated tumors, indicating a shift towards mesenchymal gene signature. The complete overall IBSP vs. CTRL differential expression analysis (DEA) (FDR<0.001, (a)), significantly enriched canonical pathways (p<0.05, (b)), upstream regulators (p<0.05, (c)), and diseases and functions associated with DEA (p<1.5e-6, (d)) can be found in **File S5**. Canonical Pathway analysis revealed a significant enrichment of FAK and integrin signaling, known regulators of glioma invasiveness. Additionally, there was also significant enrichment of Paxillin, IGF-1, and Ephrin receptor signaling which could potentially play a role in PVN-induced acquisition of a more aggressive and invasive phenotype by GBM cells (**Figure 5A**). Furthermore, IPA upstream regulator analysis predicted that NF_κ_B, STAT3 and HIF1α were significantly activated (**Figure 5C**). Based on these results, we reasoned that IBSP may induce a Mesenchymal shift in tumor cells resulting in the observed migratory phenotype described above. To further test this, we classified the CTRL and IBSP-treated cells according to TCGA classification of gliomaspheres (Laks et al., 2016) (**Figure 5C**). As expected, when we performed PCA of genes used to categorize samples of the tumors in the TCGA database (2008; Verhaak et al., 2010), these samples separated according to subclass (Classical, Proneural, Mesenchymal). We then superimposed the gene expression signatures of our control and treated gliomasphere samples. This superimposition clearly demonstrated the movement of the samples from a more Proneural state towards a more Mesenchymal signature upon IBSP treatment, particularly when comparing each CTRL to its paired IBSP sample. Though the HK157 GBM line exhibited a more robust phenotype, HK217 also displayed a similar expression profile upon IBSP treatment as its DEA (FDR<0.05) significantly overlapped with the HK157 DEA (p-value of overlap=1.3e-243).

**Figure 5:**
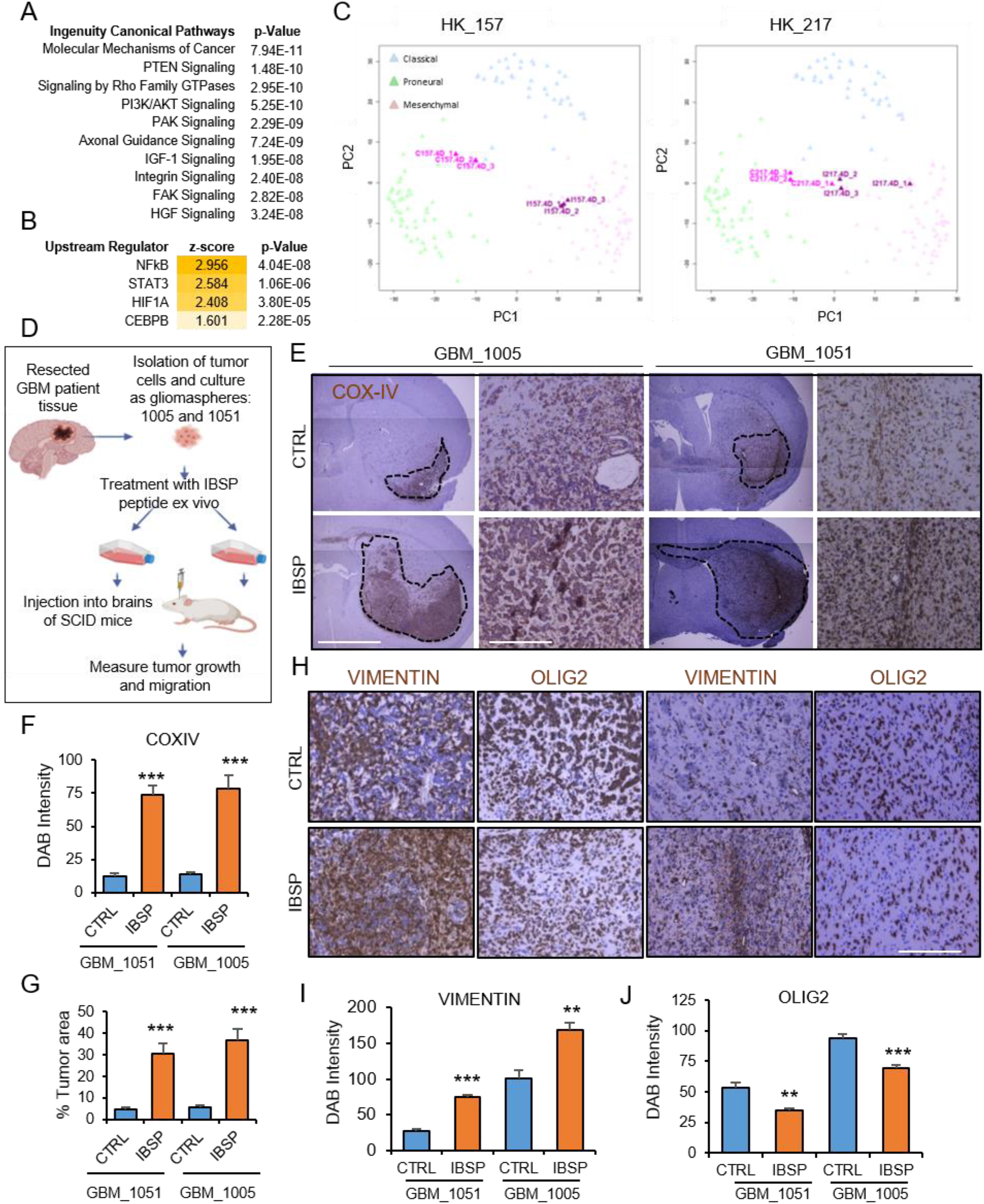
IBSP promotes a Mesenchymal gene signature in Proneural gliomaspheres. (A) Canonical pathways significantly enriched by IBSP treatment as predicted by IPA. (B) Mesenchymal pathway regulators significantly activated in IBSP-induced set of dysregulated genes (FDR<0.001) and the p<0.05. (C) PCA of GBM in the TCGA database (in background) and our HK157 (left panel) and HK217 (right panel) gliomaspheres (CTRL in pink, IBSP in purple) distributed according to gene signature used in TCGA classifications. (D) Schematic illustrates the strategy for ex vivo treatment of tumor cells with IBSP for implantation to assess tumor growth and migration (E-G) Representative images of immunohistochemistry of anti-COX IV (human mitochondrial marker) in control and IBSP treated tumors. Scale bars, 200um. Graphs show quantitation of average DAB staining intensity (COXIV staining) per group. N=3 mice, * * and * * * p<0.001 and p<0.0001, unpaired t-test. (H-J) Representative images of immunohistochemistry of mesenchymal (VIMENTIN), proneural (OLIG2) markers in control and IBSP treated tumors. Scale bars, 200um. Graphs show quantitation of average DAB staining intensity per group. N=3 mice, * * and * * * p<0.001 and p<0.0001, unpaired t-test. See also Figure S5.

To assess the behavior of IBSP-treated cells in the *in vivo* microenvironment, we pretreated two gliomasphere lines known to exhibit robust tumor formation following xenotransplantation, 1005 and 1051, for 3 days with IBSP and examined the tumors in mouse brain xenografts **(Figure 5D**). As demonstrated by staining for human cells four weeks following implantation, IBSP-treated samples produced overtly larger and more invasive tumors (**Figure 5E**), an impression substantiated by quantitative analysis of tumor area (**Figure 5F-G**). Furthermore, to assess whether IBSP potentially induced a mesenchymal shift in the tumors, we immunostained for the known Mesenchymal markers Vimentin, CD44, and YKL40 (CHI3L1), and the Proneural marker Olig2. Both the IBSP-treated tumors had significantly increased expression of the Mesenchymal markers and decreased expression of Olig2 (**Figure 5H-J and Figure S5A-B**). Our functional data thus far indicate that IBSP promotes GTC migration, proliferation of a subset of Proneural GTC, and a Mesenchymal shift in gene expression.

Since IBSP is a known integrin-binding protein, we next sought to determine which integrins, if any, might be important for its function. From the comprehensive interactome, we identified a number of expressed integrins (**Figure 6A**), some of which were previously shown to interact with the SIBLING ligands. For example, ITGαVβ3 and ITGαVβ5 interact with IBSP to promote breast cancer proliferation, and invasion, respectively (Sung et al., 1998). To determine whether the expressed integrins were relevant for IBSP function, we performed a shRNA screen based on migration capacity through the hydrogel matrix in the presence of IBSP (**Figure 6B**). Infection with at least one of the clones for ITGαV, ITGβ3, ITGα6, ITGα7, ITGβ8, and ITGα1 resulted in significantly reduced GTC dispersion in hydrogel (3≤N≤8) (**Figure 6B**). However, of all the integrins, ITGβ8 and ITGαV showed the highest expression in GTC (**Figure 6A**). IBSP is known to promote tumor invasion through its interactions with the ITGαV receptor in other cancers (Karadag et al., 2004; Sung et al., 1998). We therefore selected this subunit as the leading α-integrin candidate. Fluorescent *in situ* hybridization of patient tumor tissue revealed that ITGαV is expressed by tumor cells, including those adjacent to IBSP-expressing vasculature (**Figure S6A**). Next, we assessed its role in mediating the effects of IBSP on migration of 3 different primary gliomasphere lines representing Proneural (HK217 and HK301) and Mesenchymal (HK280) expression groups using a specific ITGαV blocking antibody. As expected, IBSP induced migration in all 3 gliomasphere lines tested (**Figure 6C**), an effect completely abolished by concomitant treatment with the ITGαV blocking antibody (**Figure 6C,D**). To validate the specificity of this effect, we knocked down ITGαV expression in tumor cells with two separate shRNAs to ITGαV (validation of knockdown is shown in **Figure S6B**,**C**). We found that ITGαV knockdown reversed the pro-migratory effects of IBSP (**Figure 6E,F**), indicating that ITGαV may act as the predominant α-integrin receptor for IBSP in GBM tumors, at least for the promigratory effect.

**Figure 6:**
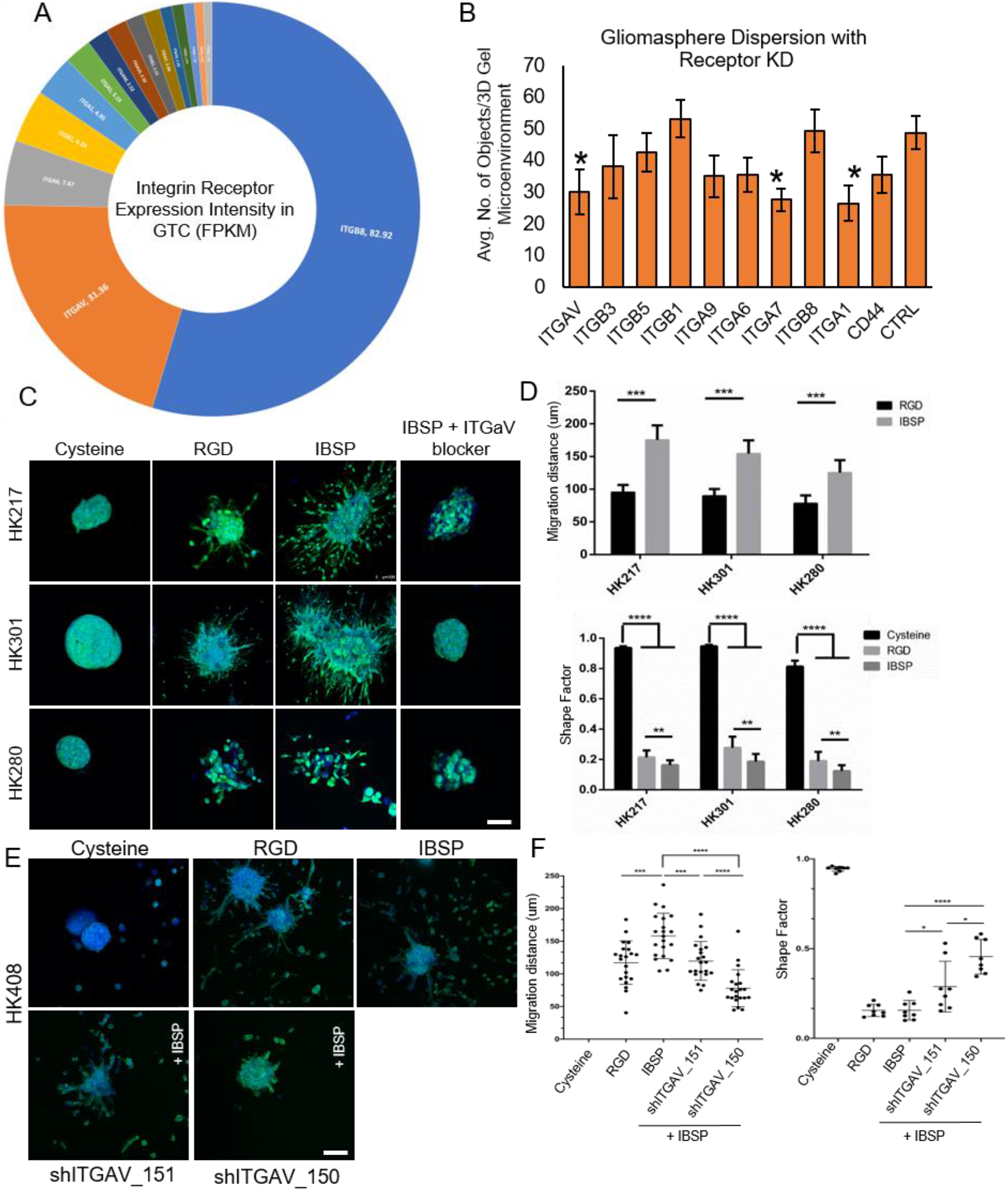
IBSP mediates its pro-migratory effects on GTC via ITGαV. (A) Pie chart shows the expression intensity of various Integrin receptors in GTC. (B) Graph shows quantitation of gliomasphere dispersion in 3D hydrogels post knockdown of integrin receptors. * p<0.05, t-test. (C,D) Representative images of GFP-labelled gliomaspheres from Proneural GBM (HK217, HK301) and Mesenchymal GBM (HK280) encapsulated in HA hydrogels and treated with RGD or IBSP peptides and IBSP+ ITGAV-neutralizing antibody. Cysteine hydrogels were used as negative control. Scale bar, 100um. Graphs show quantitation of shape factor and migration distance in RGD vs IBSP treatment groups. N=10 GBM spheroids, Mann Whitney test, * p<0.001, * * p<0.0001 (E,F) Representative images of sh-ITGAV-infected gliomaspheres treated with RGD or IBSP. Graphs show quantitation of shape factor and migration distance in RGD vs IBSP treatment groups. N=10 GBM spheroids, Mann Whitney test, * p<0.001, * * p<0.0001. See also Figure S6.

### Endogenous IBSP regulates tumor growth and migration

Our functional studies indicate that exogenously added IBSP influences GBM proliferation and migration, with the latter effect at least, requiring ITGαV. To examine the effects of endogenous, vascular-derived IBSP, we first utilized our hydrogel co-culture system. A shRNA-mediated knockdown of IBSP was performed using 3 different validated constructs in cultured primary GVC (**Figure S7A-C**). Knockdown of IBSP in GVC resulted in dramatically diminished migration of the co-cultured GTC compared to cells infected with control viruses or positive control cells (**Figure 7A-B and Figure S7D-F**). To control for any effects that IBSP expression may have had on GVC themselves, we also assessed GVC proliferation in presence of increasing concentrations of soluble IBSP, and with knockdown cells used in our studies above. Both the soluble form of IBSP and knockdown did not show any significant effect on GVC proliferation (**Figure S7G, H**), thus eliminating the likelihood of increased GVC numbers in response to our tested conditions contributing to an increase in other migration promoting factors in the co-culture system.

**Figure 7:**
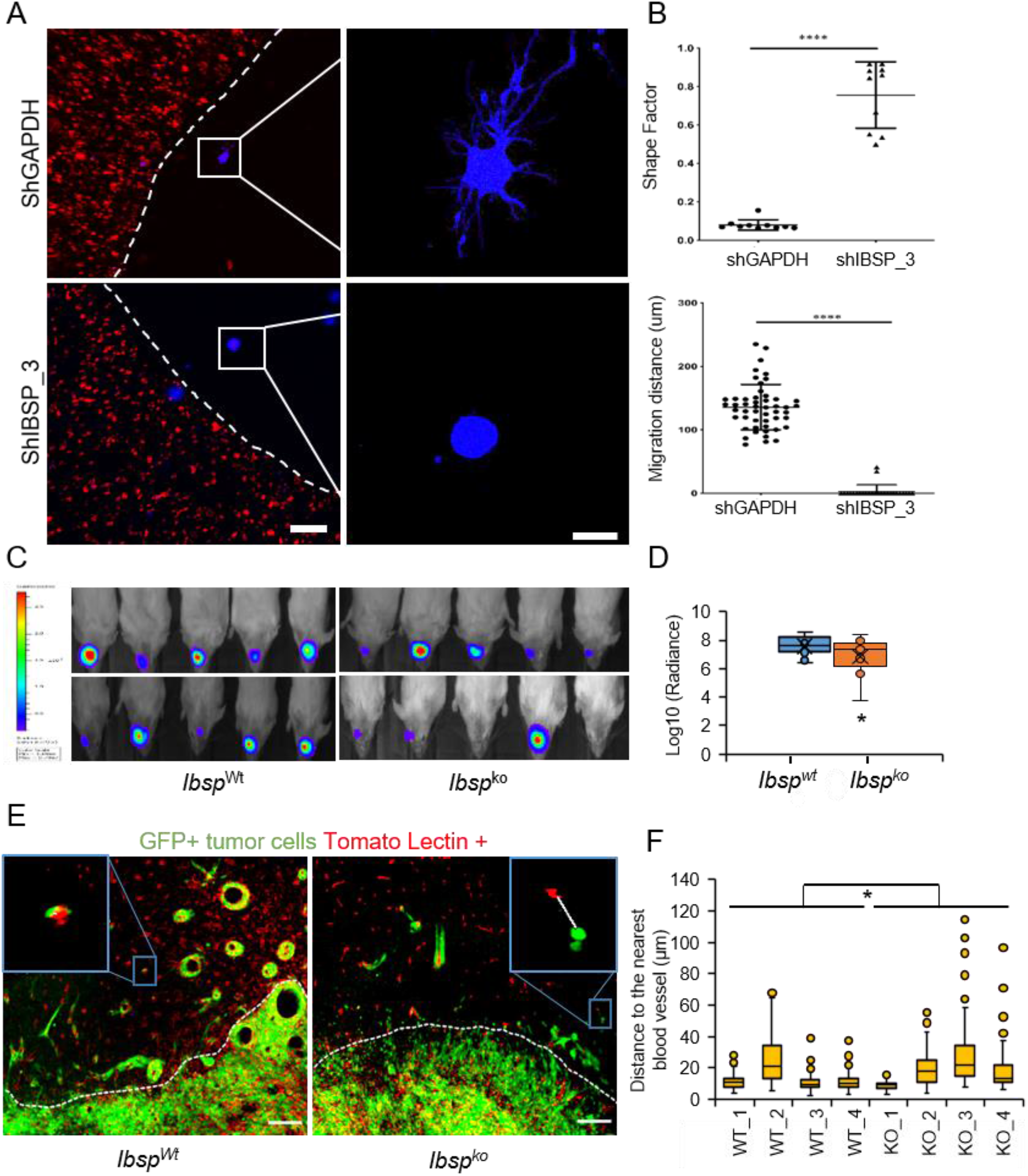
Endogenous vascular cell-derived IBSP promotes GBM growth and migration. (A,B) Representative images of hydrogels encapsulated with GBM spheroids and IBSP-deficient GVC (shIBSP) or control (shGAPDH) GVC. GVC (mCherry, red) and tumor cells (BFP, blue). White dashed lines demarcate the two hydrogels. Boxes show a higher magnification of the GBM spheroids. Scale bars, 250um. Graphs show quantitation of migration distance and shape factor in each group. N=10, One-way ANOVA and unpaired t-test, * * p<0.0001. (C,D) Images show luciferase signal from mouse GBM tumor cells implanted into IBSP knockout (Ibsp ko) and wild type (Ibsp wt) mice. Quantitation of tumor growth measured by luminescence is shown in the graph. * p<0.05, unpaired t-test, N=10 mice per group. (E) Immunostaining of tumor cells (GFP, green) and blood vessels (Tomato Lectin, red) in knockout and wild type mice. White dashed lines show the tumor edge. A representative image shown as an inlet from each group shows association of GFP+ tumor cell with Lectin+ blood vessel. Scale bars, 100um. (F) Graph shows quantitation of log normalized distance of tumor cells from blood vessel in each group using a linear mixed effects model. KO mice have higher mean distance compared to WT mice (log normalized average of 0.71, SE=0.31, p=0.02). See also Figure S7.

Next, we sought to determine whether endogenous IBSP plays a role in promoting tumor growth and migration *in vivo*. Transplantation of a murine GBM cell line expressing GFP-Luciferase into *Ibsp* WT and KO mice (Malaval et al., 2008), and quantitation of tumor growth by bioluminescence imaging revealed significantly smaller tumors (p<0.05, t-test) in *Ibsp* KO mice relative to WT controls (**Figure 7C, D**) indicating that endogenous *Ibsp* is important for GBM growth. To evaluate the potential role of *Ibsp* in migration, we examined the association of GFP-labeled tumor cells with Lectin-labeled blood vessels located outside the main body of the tumor in *Ibsp* KO and WT mice. As shown in **Figure 7E** (highlighted by dashed lines), glioma cells were more often found in close proximity to blood vessels in WT mice than in *Ibsp* KO mice, highlighting the potential role of IBSP in promoting tumor cell invasion along vessels *in vivo*. Quantifying this effect, we found that the distance between GFP+ tumor cells and lectin+ blood vessels was significantly higher in *Ibsp* KO mice than in WT mice (P=0.025, WT CI -0.57; -0.05, IBSP KO CI 0.09; 0.29) (**Figure 7F**) suggesting that *Ibsp* promotes growth and migration of tumor cells. Collectively, these findings indicate that vascular-derived IBSP plays an important role in growth and migration of GBM tumors.

## Discussion

Microvascular hyperproliferation is a major hallmark of GBM and a histopathological marker of poor prognosis (Das and Marsden, 2013). Early studies demonstrated that the vasculature can provide a trophic niche that allows for the maintenance and survival of the adjacent glioma tumor cells (Calabrese et al., 2007; Jeon et al., 2014). For example, vascular cells support GBM stem-like cells in the face of radiation treatment (Garcia-Barros et al., 2003). Another important role for the vascular niche is in supporting migration out of the main body of the tumor and into the more normal parenchyma (Gilbertson and Rich, 2007; Griveau et al., 2018) sowing the seeds for tumor recurrence and disease relapse. The molecular mechanisms mediating these effects are only beginning to be understood.

Prior studies to delineate the genes enriched in GBM vasculature compared to low grade tumors or normal brain have generally focused on the means by which the tumor induces the host vasculature to promote the genesis and ingrowth of hyperproliferative, abnormal vessels associated with GBM (Charalambous et al., 2005; Dieterich et al., 2012). The most important of these interactions include secretions of growth factors, such as VEGF and FGF, by tumor cells (Bao et al., 2006). There is ample evidence to consider the reverse signaling, where factors elaborated by the vasculature influence tumor biology. Here, we have utilized RNA sequencing and an informatics-based approach to delineate genes expressed by GBM vasculature, whose secreted extracellular factors are predicted to interact with proteins expressed by tumor cells. Our study is meant to provide a broader picture of the perivascular niche-tumor interactome in order that other mechanisms can be targeted.

A key question to be asked is: which interactome is most significant, one with a focus on those molecules enriched in both GBM vasculature and GBM tumor cells or one that is broader, taking into account all potential interactions? From a therapeutic perspective, there would be an inherent desirability to target pathways and factors that are enriched in GBM compared to normal brain. However, when considering the complete biology of the vascular-tumor interaction, proteins that are expressed by the glioma vasculature and the tumor cells, but not necessarily enriched compared to normal cells will still have the potential to play important roles in tumor progression. For this reason, we have described a more global interactome in addition to ones including only glioma-enriched genes.

One potential limitation of our study is in the choice of tumor and vascular cell populations. We selected the A2B5-expressing population because it has been previously demonstrated to encompass the glioma tumor-initiating cells (Auvergne et al., 2013; Ogden et al., 2008). Furthermore, this population can be compared directly to non-cancerous A2B5+ glial progenitors, as has been done previously (Auvergne et al., 2013). In line with these studies, we found that this fraction expresses numerous genes in common with normal glial progenitors, but that it also has dysregulated oncogenic pathways, indicating that they are true cancer cells. However, it is possible that we have missed key tumor cells in our analysis that may not express A2B5. Similarly, we used CD31 to highly identify GVC. However, it is likely that other cells that are tightly associated with endothelial cells, including pericytes and perivascular immune cells, may be present within this fraction. Rather than considering such cells as “contaminants”, their lack of exclusion allows for the consideration of the vascular niche as a whole rather than on a cell-by-cell basis.

Our informatics analysis provides strong support for the concept that a major role for the GBM vasculature is in the promotion of tumor cell migration. Pro-migratory functions dominated the gene ontologies of the interactome. *In vitro* analysis using a novel hydrogel-based co-culture system supports this hypothesis, as short-term, primary GVC produce robust migration of cells out of GBM spheroid cultures. Our findings, however, do not preclude other roles for GVC-derived factors. Such roles could include effects on tumor cell survival and proliferation, with the latter function demonstrated for IBSP in this study. In addition, GVC-derived factors could play important roles and influence other cells in the tumor microenvironment, including immune cells.

As a proof of concept, we examined a relatively poorly studied protein, IBSP, and demonstrated its strong pro-migratory effects *in vitro* and *in vivo*. Given that there are numerous integrin-binding proteins associated with the tumor microenvironment, including other members of the SIBLING family, especially OPN (Lamour et al., 2015), we were surprised by the significance of IBSP, as knockdown of IBSP in GVC appeared to completely abolish migration of tumor cells in our culture system, and tumors implanted into *Ibsp* ko mice showed a significantly reduced number of vascular-associated tumor cells outside the body of the tumor. However, these findings do not imply that IBSP is the only factor that regulates this migration, as it is likely that IBSP acts in concert with other factors. The mechanisms by which IBSP promotes GBM migration remain to be fully elucidated, although it is clear that IBSP requires interaction with ITGαV for its pro-migration effect.

A somewhat surprising result was that in addition to promoting migration, IBSP also promoted the proliferation of GBM cells, but this effect, at least *in vitro*, was associated only with the Proneural GBM lines tested. It is striking to note that IBSP expression is only associated with worse outcome in Proneural tumors. The reasons why the *in vitro* pro-proliferative effects of IBSP are restricted to Proneural cells are unknown. Cells of all three tumor subtypes express ITGαV and have a pro-migratory response to IBSP and, thus, it is not simply a matter of cells being insensitive to IBSP signaling. It is possible that the pro-proliferative response is due to action at other receptors or the activation of downstream pathways in Proneural cells that are not activated in cells of the other 2 main subtypes, but the resolution of this question will require further study.

In conclusion, our study utilized RNA sequencing, coupled with bioinformatics approaches, to describe a comprehensive GVC-GTC interactome. While we anticipate that the IBSP-ITGαV axis will be considered a target for therapeutic intervention, our elucidation of the broader GVC-GTC interactome is meant to serve as a resource for the brain tumor and, indeed, the cancer research community at large.

## Experimental procedures

See supplemental information

## Acknowledgements

The authors thank the UCLA BTTR and Dr. William Yong, the UCLA Neuroscience Genomics Core for RNA amplification and cDNA library preparation, the Jonsson Comprehensive Cancer Center Flow Cytometry Core, the UCLA Broad Stem Cell Research Center for RNA sequencing, Dr. Xinmin Li and the UCLA Clinical Microarray Core in the Department of Pathology for technical contributions. We thank Dr. Robert Damoiseaux and the UCLA Molecular Screening Shared Resource for invaluable assistance with our limited shRNA screen. We thank Holly Wilhalme in the UCLA Institute for Digital Research and Education (IDRE) for aid in figure presentation. The IBSP null mice were generously supplied by Dr. Martha Sommerman and Dr. Harvey Goldberg at the National Institute of Arthritis and Musculoskeletal and Skin Diseases. The IDH WT murine glioma line was a kind gift from Drs. Maria Castro and Pedro Lowenstein.

## Grant support

This work was supported by grants from the Dr. Miriam and Sheldon G. Adelson Medical Research Foundation, The National Institutes of Health Grants P50 CA211015, CA241927-01A1 and CA197943. This work was also supported by a University of California Cancer Research Coordinating Committee Award, Broad Stem Cell Research Center Seed Grant, and an American Brain Tumor Association Discovery Award.

## Author contributions

Conceptualization: YG, AS, SDM, LIA, IN, GC, SAG, STC, SLS, HIK. Methodology: SKS, LIA, GC, RK. Formal Analysis YG, AS, SDM, MC, FG, YQ, RK. Investigation YG, AS, SDM, RK, MC, SB, JM, NR, DRL. Resources LML, GWM, SKS, HIK. Writing - Original Draft: YG, AS, SDM, HIK. Supervision: LIA, GC, SKS, IN, HIK.

## Declaration of Interests

The authors declare no competing interests.

## Notes

### Competing Interest Statement

The authors have declared no competing interest.

## References

Auvergne, R.M., Sim, F.J., Wang, S., Chandler-Militello, D., Burch, J., Al Fanek, Y., Davis, D., Benraiss, A., Walter, K., Achanta, P., et al. (2013). Transcriptional differences between normal and glioma-derived glial progenitor cells identify a core set of dysregulated genes. Cell Rep 3, 2127–2141.

Bao, S., Wu, Q., Sathornsumetee, S., Hao, Y., Li, Z., Hjelmeland, A.B., Shi, Q., McLendon, R.E., Bigner, D.D., and Rich, J.N. (2006). Stem cell-like glioma cells promote tumor angiogenesis through vascular endothelial growth factor. Cancer Res 66, 7843–7848.

Bellahcène, A., Castronovo, V., Ogbureke, K.U.E., Fisher, L.W., and Fedarko, N.S. (2008). Small integrin-binding ligand N-linked glycoproteins (SIBLINGs): multifunctional proteins in cancer. Nat Rev Cancer 8, 212–226.

Bowman, R.L., Wang, Q., Carro, A., Verhaak, R.G.W., and Squatrito, M. (2017). GlioVis data portal for visualization and analysis of brain tumor expression datasets. Neuro Oncol 19, 139–141.

Brat, D.J., and Van Meir, E.G. (2001). Glomeruloid microvascular proliferation orchestrated by VPF/VEGF: a new world of angiogenesis research. Am J Pathol 158, 789–796.

Brooks, L.J., and Parrinello, S. (2017). Vascular regulation of glioma stem-like cells: a balancing act. Curr Opin Neurobiol 47, 8–15.

Calabrese, C., Poppleton, H., Kocak, M., Hogg, T.L., Fuller, C., Hamner, B., Oh, E.Y., Gaber, M.W., Finklestein, D., Allen, M., et al. (2007). A perivascular niche for brain tumor stem cells. Cancer Cell 11, 69–82.

Charalambous, C., Hofman, F.M., and Chen, T.C. (2005). Functional and phenotypic differences between glioblastoma multiforme-derived and normal human brain endothelial cells. J Neurosurg 102, 699–705.

Charalambous, C., Virrey, J., Kardosh, A., Jabbour, M.N., Qazi-Abdullah, L., Pen, L., Zidovetzki, R., Schönthal, A.H., Chen, T.C., and Hofman, F.M. (2007). Glioma-associated endothelial cells show evidence of replicative senescence. Exp Cell Res 313, 1192–1202.

Charles, N., and Holland, E.C. (2010). The perivascular niche microenvironment in brain tumor progression. Cell Cycle 9, 3012–3021.

Charles, N., Ozawa, T., Squatrito, M., Bleau, A.-M., Brennan, C.W., Hambardzumyan, D., and Holland, E.C. (2010). Perivascular nitric oxide activates notch signaling and promotes stem-like character in PDGF-induced glioma cells. Cell Stem Cell 6, 141–152.

Dabney, A.R. (2006). ClaNC: point-and-click software for classifying microarrays to nearest centroids. Bioinformatics 22, 122–123.

Das, S., and Marsden, P.A. (2013). Angiogenesis in glioblastoma. N. Engl. J. Med. 369, 1561–1563.

Dieterich, L.C., Mellberg, S., Langenkamp, E., Zhang, L., Zieba, A., Salomäki, H., Teichert, M., Huang, H., Edqvist, P.-H., Kraus, T., et al. (2012). Transcriptional profiling of human glioblastoma vessels indicates a key role of VEGF-A and TGFβ2 in vascular abnormalization. J Pathol 228, 378–390.

Farin, A., Suzuki, S.O., Weiker, M., Goldman, J.E., Bruce, J.N., and Canoll, P. (2006). Transplanted glioma cells migrate and proliferate on host brain vasculature: a dynamic analysis. Glia 53, 799–808.

Garcia-Barros, M., Paris, F., Cordon-Cardo, C., Lyden, D., Rafii, S., Haimovitz-Friedman, A., Fuks, Z., and Kolesnick, R. (2003). Tumor response to radiotherapy regulated by endothelial cell apoptosis. Science 300, 1155–1159.

Gilbertson, R.J., and Rich, J.N. (2007). Making a tumour’s bed: glioblastoma stem cells and the vascular niche. Nat Rev Cancer 7, 733–736.

Griveau, A., Seano, G., Shelton, S.J., Kupp, R., Jahangiri, A., Obernier, K., Krishnan, S., Lindberg, O.R., Yuen, T.J., Tien, A.-C., et al. (2018). A Glial Signature and Wnt7 Signaling Regulate Glioma-Vascular Interactions and Tumor Microenvironment. Cancer Cell 33, 874-889.e7.

Hemmati, H.D., Nakano, I., Lazareff, J.A., Masterman-Smith, M., Geschwind, D.H., Bronner-Fraser, M., and Kornblum, H.I. (2003). Cancerous stem cells can arise from pediatric brain tumors. Proc Natl Acad Sci U S A 100, 15178–15183.

Jeon, H.-M., Kim, S.-H., Jin, X., Park, J.B., Kim, S.H., Joshi, K., Nakano, I., and Kim, H. (2014). Crosstalk between glioma-initiating cells and endothelial cells drives tumor progression. Cancer Res 74, 4482–4492.

Johnson, W.E., Li, C., and Rabinovic, A. (2007). Adjusting batch effects in microarray expression data using empirical Bayes methods. Biostatistics 8, 118–127.

Karadag, A., Ogbureke, K.U.E., Fedarko, N.S., and Fisher, L.W. (2004). Bone sialoprotein, matrix metalloproteinase 2, and alpha(v)beta3 integrin in osteotropic cancer cell invasion. J Natl Cancer Inst 96, 956–965.

Krishnan, S., Szabo, E., Burghardt, I., Frei, K., Tabatabai, G., and Weller, M. (2015). Modulation of cerebral endothelial cell function by TGF-β in glioblastoma: VEGF-dependent angiogenesis versus endothelial mesenchymal transition. Oncotarget 6, 22480–22495.

Laks, D.R., Masterman-Smith, M., Visnyei, K., Angenieux, B., Orozco, N.M., Foran, I., Yong, W.H., Vinters, H.V., Liau, L.M., Lazareff, J.A., et al. (2009). Neurosphere formation is an independent predictor of clinical outcome in malignant glioma. Stem Cells 27, 980–987.

Laks, D.R., Crisman, T.J., Shih, M.Y.S., Mottahedeh, J., Gao, F., Sperry, J., Garrett, M.C., Yong, W.H., Cloughesy, T.F., Liau, L.M., et al. (2016). Large-scale assessment of the gliomasphere model system. Neuro Oncol 18, 1367–1378.

Lamour, V., Henry, A., Kroonen, J., Nokin, M.-J., von Marschall, Z., Fisher, L.W., Chau, T.-L., Chariot, A., Sanson, M., Delattre, J.-Y., et al. (2015). Targeting osteopontin suppresses glioblastoma stem-like cell character and tumorigenicity in vivo. Int J Cancer 137, 1047–1057.

Malaval, L., Wade-Guéye, N.M., Boudiffa, M., Fei, J., Zirngibl, R., Chen, F., Laroche, N., Roux, J.-P., Burt-Pichat, B., Duboeuf, F., et al. (2008). Bone sialoprotein plays a functional role in bone formation and osteoclastogenesis. J Exp Med 205, 1145–1153.

Musumeci, G., Castorina, A., Magro, G., Cardile, V., Castorina, S., and Ribatti, D. (2015). Enhanced expression of CD31/platelet endothelial cell adhesion molecule 1 (PECAM1) correlates with hypoxia inducible factor-1 alpha (HIF-1α) in human glioblastoma multiforme. Exp Cell Res 339, 407–416.

Nunez, F.J., Mendez, F.M., Kadiyala, P., Alghamri, M.S., Savelieff, M.G., Garcia-Fabiani, M.B., Haase, S., Koschmann, C., Calinescu, A.-A., Kamran, N., et al. (2019). IDH1-R132H acts as a tumor suppressor in glioma via epigenetic up-regulation of the DNA damage response. Sci Transl Med 11.

Ogden, A.T., Waziri, A.E., Lochhead, R.A., Fusco, D., Lopez, K., Ellis, J.A., Kang, J., Assanah, M., McKhann, G.M., Sisti, M.B., et al. (2008). Identification of A2B5+CD133-tumor-initiating cells in adult human gliomas. Neurosurgery 62, 505–514; discussion 514-515.

Pen, A., Moreno, M.J., Martin, J., and Stanimirovic, D.B. (2007). Molecular markers of extracellular matrix remodeling in glioblastoma vessels: microarray study of laser-captured glioblastoma vessels. Glia 55, 559–572.

Scholz, A., Harter, P.N., Cremer, S., Yalcin, B.H., Gurnik, S., Yamaji, M., Di Tacchio, M., Sommer, K., Baumgarten, P., Bähr, O., et al. (2016). Endothelial cell-derived angiopoietin-2 is a therapeutic target in treatment-naive and bevacizumab-resistant glioblastoma. EMBO Mol Med 8, 39–57.

Smyth, G.K., Michaud, J., and Scott, H.S. (2005). Use of within-array replicate spots for assessing differential expression in microarray experiments. Bioinformatics 21, 2067–2075.

Stratmann, A., Risau, W., and Plate, K.H. (1998). Cell type-specific expression of angiopoietin-1 and angiopoietin-2 suggests a role in glioblastoma angiogenesis. Am J Pathol 153, 1459–1466.

Sung, V., Stubbs, J.T., Fisher, L., Aaron, A.D., and Thompson, E.W. (1998). Bone sialoprotein supports breast cancer cell adhesion proliferation and migration through differential usage of the alpha(v)beta3 and alpha(v)beta5 integrins. J Cell Physiol 176, 482–494.

Tchoghandjian, A., Baeza, N., Colin, C., Cayre, M., Metellus, P., Beclin, C., Ouafik, L., and Figarella-Branger, D. (2010). A2B5 cells from human glioblastoma have cancer stem cell properties. Brain Pathol 20, 211–221.

Verhaak, R.G.W., Hoadley, K.A., Purdom, E., Wang, V., Qi, Y., Wilkerson, M.D., Miller, C.R., Ding, L., Golub, T., Mesirov, J.P., et al. (2010). Integrated genomic analysis identifies clinically relevant subtypes of glioblastoma characterized by abnormalities in PDGFRA, IDH1, EGFR, and NF1. Cancer Cell 17, 98–110.

Xiao, W., Zhang, R., Sohrabi, A., Ehsanipour, A., Sun, S., Liang, J., Walthers, C.M., Ta, L., Nathanson, D.A., and Seidlits, S.K. (2018). Brain-Mimetic 3D Culture Platforms Allow Investigation of Cooperative Effects of Extracellular Matrix Features on Therapeutic Resistance in Glioblastoma. Cancer Res 78, 1358–1370.

Yan, G.-N., Yang, L., Lv, Y.-F., Shi, Y., Shen, L.-L., Yao, X.-H., Guo, Q.-N., Zhang, P., Cui, Y.-H., Zhang, X., et al. (2014). Endothelial cells promote stem-like phenotype of glioma cells through activating the Hedgehog pathway. J Pathol 234, 11–22.

