## Supplementary information for "Generation of a molecular interactome of the glioblastoma perivascular niche reveals Integrin Binding Sialoprotein as a key mediator of tumor cell migration"

**Supplemental Information**

**Supplementary Tables:**

**Table S1: Patient Information**
(A) Non-transformed samples, Primary Brain Glial Progenitor Cells (BGPC), Primary Brain Vascular cells (BVC), Grey Matter (GM), White Matter (WM).

| Sample # | Cell Fraction Isolated for RNA-seq | Age | Gender | Surgical notes: |
|---|---|---|---|---|
| 12 | BVC, BGPC | 19 | M | Left hemispherectomy for Left MCA perinatal stroke |
| 13 | BVC, BGPC | 13 | M | Left temporal occipital craniotomy for cortical dysplasia |
| 14 | GM and WM, BVC | 5 | F | Left frontal temporal craniotomy for resection of left cortical dysplasia. |
| 15 | GM and WM | 12 | F | Right hemispherectomy for diffuse right hemisphere cortical malformation and seizures with Lennox-Gastaut. |
| 16 | GM and WM | 3 | F | Right hemicraniotomy for epilepsy secondary to right intraventricular teratoma. |

(B) GBM Tumor samples GBM Tumor Cells (GTC), GBM Vascular Cells (GVC).

| Sample # | Diagnosis | Cell Fraction Isolated for RNA-seq | Age | Gender | Characteristics and Cytogenetics |
|---|---|---|---|---|---|
| GBM4 | GBM Primary with gigantocellular features | Unsorted, GVC, GTC | 66 | M | Maximum Ki67: 90%; MGMT Not Methylated; Multiple copies 7, 1,19. Monosomy 19. 10q loss |
| GBM5 | GBM, Primary with oligodendroglial component | Unsorted, GVC | 42 | M | Maximum Ki67: 70%; MGMT Methylated; IDH1 positive; Monosomy 10 |
| GBM6 | GBM Primary | Unsorted, GVC | 77 | F | Maximum Ki67: 40%; MGMT Methylated; EGFR amplified |
| GBM7 | GBM Primary | Unsorted, GVC, GTC | 55 | F | Maximum Ki67: 20%; MGMT Not Methylated; EGFR amplified; Monosomy 10 |
| GBM8 | GBM Primary | Unsorted, GVC, GTC | 36 | M | Maximum Ki67: 40%; EGFR amplified; Monosomy 10 |
| GBM9 | GBM Primary | Unsorted, GVC, GTC | 59 | M | Maximum Ki67: 40%; MGMT Not Methylated; EGFR amplified; EGFR vIII positive |
| GBM10 | GBM Primary (outside report suggests gliosarcoma) | Unsorted, GVC, GTC | 58 | M | Maximum Ki67: 70%; MGMT Not Methylated; Polysomy 1 and 19. 10q loss |

**Table S2: Key resources**

| Reagents | Source | Identifier |
|---|---|---|
| **Antibodies** | | |
| Mouse monoclonal anti- CD31, JC70A, human | Agilent DAKO | M082329-2 |
| Anti-human CD31 antibody, REA730 | Miltenyi Biotec | 130-110-670 |
| Rabbit polyclonal anti-GFP | Novus Biologicals | NB600-308 |
| Mouse monoclonal anti-mCherry | Novus Biologicals | NBP1-96752 |
| Chicken polyclonal anti-GFP | Novus Biologicals | NB100-1614 |
| Chicken polyclonal anti-mCherry | Millipore Sigma | AB356481 |
| Tomato Lectin-DyLight 649 | Vector Laboratories | DL-1178-1 |
| Anti-A2B5 antibody, clone 105 | ATCC | |
| IBSP antibody, LFMB-25 | Santa Cruz Biotechnology | |
| ITGαV antibody | Abcam | ab16821 |
| Anti-COXIV | Cell Signaling | 4850 |
| Anti-VIMENTIN | Cell Signaling | 5741 |
| Anti-OLIG2 | Abcam | Ab109186 |
| Anti-CD44 | Cell Signaling | 3570 |
| Anti-YKL40/CHI3L1 | Abcam | Ab77528 |
| **Chemicals/commercial assays** | | |
| Percoll Plus | GE healthcare | |
| Collagenase II and IV | Worthington Biochemical | LS004174, LS004186 |
| Hibernate A minus Ca, Mg | BrainBits LLC | HACAMG500 |
| Endothelial cell media | Sciencell | 1001 |
| Endothelial cell media (EGM-2) | R&D systems | CCM207 |
| DMEM:F12 | Invitrogen | |
| Anti-human CD31 Dynabeads | Thermofisher Scientific | |
| RGD peptide | Genscript | |
| Recombinant Bone Sialoprotein (IBSP) | Abcam | ab219248 |
| Recombinant Lumican | Abcam | ab114635 |
| Recombinant InhibinB-A | Abcam | ab53506 |
| Recombinant WNT5A | Abcam | ab204627 |
| Vectastain Elite ABC-HRP, DAB Kit | Vector Laboratories | PK6100, and SK4100 |
| TSA Plus Cyanine 3 System | Perkin Elmer | NEL744001KT |
| TSA Plus Cyanine 5 Evaluation Kit | Perkin Elmer | NEL745E001KT |
| DiI-Ac-LDL assay | Cell applications | 022k |
| CCK8 assay | Dojindo Molecular Technologies | CK04 |
| **Experimental Models: Cell Lines** | | |
| Patient-derived gliomasphere lines | This lab | N/A |
| Human Brain microvascular endothelial cells (HBEC/BVC) | Sciencell | 1000 |
| 293T packaging cells | ATCC | CRL-11268 |

| | | |
|---|---|---|
| Murine GBM line | Laboratory of Dr. Maria Castro | Nunez et al., 2019 |
| **Mouse strains** | | |
| NOD.Cg-$Prkdc^{scid}$ $Il2rg^{tm1Wjl}$/SzJ (NSG) | Jackson Laboratory | 00557 |
| C57BL6 | Jackson Laboratory | 000664 |
| IBSP knockout mice ($Ibsp^{tm1Jeau}$/$Ibsp^{tm1Jeau}$), 129/SvJ-CD1 background | - | Martha Sommerman and Harvey Goldberg labs |
| **Lentiviral vectors** | | |
| pLV-FireflyLuc-GFP | UCLA Vector Core | |
| pLV(EXP)-puro-EF1A-mCherry | Vector Builder | |
| pGIPZ_puro_shRNA_IBSP | Horizon Discovery | VHG5518, clones V2LHS_61769, V3LHS_334208, V3LHS_334212 |
| pGIPZ_puro_shRNA_ITGaV | Horizon Discovery | VHG5518, clones V2LHS_133468, V3LHS_365150, V3HLS_365151 |

**Table S3: Primers**

| Genes | Forward primer | Reverse Primer |
|---|---|---|
| **Human 18srRNA** | GGCCCTGTAATTGGAATGAGTC | CCAAGATCCAACTACGAGCTT |
| **Human IBSP** | CACTGGAGCCAATGCAGAAGA | TGGTGGGGTTGTAGGTTCAAA |
| **Human ITGαV** | ATCTGTGAGGTCGAAACAGGA | ATCTGTGAGGTCGAAACAGGA |

**Experimental procedures:**

**Human Tissue Samples:** Human brain tissue samples were obtained from surgical resections from patients undergoing GBM tumor resection surgeries or brain surgeries for treatment of non-neoplastic conditions (non-transformed cortical resections) under an approved University of California, Los Angeles (UCLA) Institutional Review Board (IRB) protocol. Information on patient diagnoses and sample characteristics can be found in Tables S1A and S1B. Our non-transformed samples were collected from children and adolescents (3-19 years old) and consist of small sections of healthy cortex or white matter, which were resected to gain access to deeper epileptic or otherwise pathological brain structures, and were considered normal according to MRI, and electroencephalogram studies (table S1A). In total, we sequenced 8 primary GBM samples with no prior treatments, their GVC cell fractions, and from 5 of which we isolated the GTC cellular fractions (Supplementary Table 1B). Samples from which specific cell types were isolated, were collected either in the operating room (non-transformed) or immediately following surgery through the pathologist, which allowed for a limited time (8-10 hours) between tissue resection and our purification schemes (Figure 2A). RNA extraction immediately followed cell type isolation, thus best representing the *in vivo* transcriptome of the cells, as acquired through whole-genome RNA sequencing (RNA-seq). RNA-seq was also carried out on small pieces of un-dissociated parent GBMs and non-transformed grey matter (GM) and white matter (WM) as controls. Un-dissociated whole samples, i.e. GBM, and GM/WM, were stored in RNAlater (Thermo Fisher Scientific) at -20ºC, or flash frozen immediately following resection, respectively. The un-dissociated samples were not used in downstream analyses and were only included as quality check controls. Brain tumor samples were collected in collaboration with the UCLA Brain Tumor Translational Resource (BTTR) and were graded by the attending neuropathologist according to guidelines set forth by the World Health Organization (WHO). All tissues were collected with informed patient/guardian consent.

**Tissue processing:** To obtain a single cell suspension from the resected tissues, samples (300-700 mg) were minced into ~1mm pieces, and incubated in 12500 U of Collagenase II and Collagenase IV in Hibernate A media at 37ºC for 20 min with gentle agitation every 5min. Following enzymatic digestion, the cell suspension was passed through a 100µm filter and cells were pelleted at 1000xg for 5min. To remove cellular debris and red blood cells (RBC), the cell pellet was re-suspended in a total volume of 1.5ml of DMEM/F12, and 1.5ml of working 1X Percoll solution. RBCs were pelleted by centrifugation of this suspension at 1000xg for 5min. To the remaining supernatant and debris 1.5ml of 4X buffer was gradually added, to facilitate a shift in osmolality that would selectively allow live cells to pellet and centrifuged at 3000xg for 7min. Cells in the pellet were gradually re-introduced to normal salt concentrations by addition of 10-15ml media, were passed through a 40µm filter, and were again pelleted by centrifugation at

1000xg for 5min. The cells were then re-suspended in 1ml of phosphate buffered saline (PBS) containing 0.1% Bovine Serum Albumin and the number of cells and their viability were assessed.

**Cell-type enrichment and purification by MACS and FACS:** A vascular cell enrichment protocol was carried out using anti-human CD31 (PECAM-1) according to the manufacturer's protocol for endothelial cell positive selection. From the EC-depleted fraction, A2B5+ cells were then isolated by FACS, as previously described (Auvergne et al., 2013). Briefly, cells were incubated in 500µl of A2B5 antibody supernatant for 30 minutes at 4ºC. The cells were washed in 10ml of PBS and incubated in Alexa Fluor 488 Goat anti-mouse IgM secondary antibody (1:1000 in PBS with 0.5% BSA) for 30 minutes at 4ºC. The cells were washed in 10ml of PBS, re-suspended at 1 million/ml in Hibernate A supplemented with B27 for dead cell exclusion, and 5µM DRAQ7 for nucleated cell inclusion. Appropriate isotype controls and un-stained cells were included, and cells were sorted on a FACS ARIA flow cytometer. Immediately following EC and A2B5+ cell preparation protocols, cells were lysed in 1ml QIAzol lysis reagent. RNA isolation was carried out according to manufacturer's protocol RNA concentration and quality were assessed by NanoDrop spectrophotometer.

**RNA Sequencing and analysis:** Total RNA integrity was examined using the Agilent Bioanalyzer 2000. 100ng of cDNA were used in the library preparation All samples were multiplexed into a single pool in order to avoid batch effects and sequenced using an Illumina HiSeq 2000 sequencer. 45 million reads per sample were obtained, and quality control was performed on base qualities and nucleotide composition of sequences. Alignment to the H.sapiens (Hg38) refSeq reference gene annotation was performed using the STAR spliced read aligner with default parameters. Between 60 and 82% (avg 76%) of the reads mapped uniquely to the human genome. Total counts of read-fragments aligned to candidate gene regions were derived using HTSeq program (www.huber.embl.de/users/anders/HTSeq/doc/overview.html) with Human Hg38 refSeq as a reference and used as a basis for the quantification of gene expression. Only uniquely mapped reads were used for subsequent analyses. Differential expression analysis was conducted with R-project and the Bioconductor package edgeR. Statistical significance of the differential expression, expressed as $Log_2$ Fold Change (logFC), was determined, using tag-wise dispersion estimation, at p-Value of <0.005 unless stated otherwise. FPKM values were reported as measure of relative expression units.

**Ingenuity pathway analysis:** IPA (www.ingenuity.com, QIAGEN) was used in determining cellular localization of GTC/EC genes, and identifying direct and indirect protein interactions among EC extracellular factors and GTC PM molecules, along with manual curation and minimal use of the STRING functional protein association networks online tool (http://string-db.org), which was instrumental in developing both interactomes (according to DEA, and the comprehensive interactome according to FPKM expression units). Canonical pathways, upstream regulators and disease and functions associated with a gene list were considered to be significant at p<0.05 unless state otherwise.

**Gliomasphere cultures**: GBM-derived primary human cultures were previously described (Hemmati et al., 2003; Laks et al., 2009). Gliomaspheres (neurospheres) were dissociated down to single cells with Accumax (Sigma) every 7-14 days depending on growth rate.

**Immunohistochemistry:** Formalin fixed paraffin embedded (FFPE) patient GBM tissue blocks and tumor sections from mouse xenografts were sectioned at a thickness at 10µm and 5um respectively. Antigen retrieval was performed on deparaffinized and rehydrated sections with 0.1M sodium citrate, pH 6.1 and pepsin-mediated antigen retrieval. Endogenous peroxidase activity was quenched in 0.3% $H_2O_2$ in TBS. Sections were incubated in blocking solution (5% Normal Goat or Donkey serum and 1% BSA in TBST). Appropriate primary antibodies were added and incubated overnight at 4ºC in a humidified chamber. Biotinylated secondary antibody, and HRP-antibody were added and incubated at RT. DAB kit was used to visualize the antibody staining and hematoxylin was used as nuclear counter stain.

**Microarray-based gene expression analysis:** Concentration and quality of RNA samples were examined using the NanoDrop ND-1000 Spectrophotometer (NanoDrop Technologies) and the Agilent 2100 Bioanalyzer (Agilent Technologies). RNA samples were reverse transcribed and labeled according to the manufacturer's instructions and hybridized to Affymetrix high-density oligonucleotide HG-U133A Plus 2.0 Human Arrays. Microarray data analysis was performed as described previously. Briefly, array preprocessing was completed in the R computing environment (http://www.r-project. org) using Bioconductor packages (http://www.bioconductor.org). Raw data were normalized using the robust multiarray method (12582260). To eliminate batch effects, additional normalization was performed using the R package "ComBat" (http://statistics.byu.edu/johnson/ComBat; 16632515) with default parameters. Contrast analysis of differential expression was performed using the LIMMA package. After linear model fitting, a Bayesian estimate of differential expression was calculated using a modified t test. The threshold for statistical significance was set at $P < 0.005$ for differential expression analysis and $P < 0.01$ for explorative analyses (gene ontology and pathway analysis). Gene ontology and pathway analysis were carried out using the Database for Annotation, Visualization and Integrated Discovery (DAVID, https://david.ncifcrf.gov), GSEA (16199517), and Ingenuity Pathway Analysis (IPA; www.ingenuity.com).

**TCGA gliomasphere classification:** Our gliomasphere transcriptomes were classified into the three (Classical, Mes, and PN) clinically relevant TCGA sub-classifications as previous

described (Laks et al., 2016). Briefly, the 173 core TCGA glioblastoma samples used in TCGA subclassifications of GBMs (Verhaak et al., 2010) were used to build our classification model. The TCGA unified gene expression data (across three microarray platforms: Affymetrix HuEx array, Affymetrix U133A array and Agilent 244K) were combined with our gliomasphere data from Affymetrix U133 plus 2.0 array, and utilizing the limma R package, they were normalized together (Smyth et al., 2005). Following batch effect normalization using the ComBat R package (http://statistics.byu.edu/johnson/ComBat/)
(Johnson et al., 2007), we used the LDA based centroid classification algorism (ClaNC) used by (Verhaak et al., 2010) to develop a 3-class centroid-based classifier from 38 Classical, 56 Mes, and 53 PN TCGA samples (Dabney, 2006), where the 26 TCGA neural samples were excluded. Only 789 of the of the 840 TCGA classifier genes were used to assign classifications to our gliomaspheres, due to limitations in gene name overlap between TCGA and our platforms.

**HA Thiolation:** High molecular weight hyaluronic acid (HA) was thiolated according to established protocols (Xiao et al., 2018). In all cases, molar ratios are reported with respect to HA carboxyl groups. Briefly, sodium hyaluronate (500-750 kDa, $M_W$ = 700 kDa, Life core) was dissolved in deionized water (DI-$H_2O$). Next, 1-ethyl-3-[3-dimethylaminopropyl]carbodiimide (EDC, Thermofisher Scientific) was dissolved in DI-$H_2O$ and added to the solution at a 0.25 molar ratio. N-hydroxysuccinimide (NHS, Acros Organics) was then added to the HA solution at a 0.125 molar ratio. The solution beaker was stirred continuously at room temperature (RT) while pH was adjusted to 5.50 using 1 M HCl for 45 minutes. Then, cystamine dihydrochloride (Sigma-Aldrich) was added to the reaction at a molar ratio of 0.25 and pH was adjusted to 6.25 using 1 M NaOH. The reaction was continuously stirred at RT overnight. The next day, dithiothreitol (DTT, Sigma-Aldrich) was added (1 molar ratio), the solution pH adjusted to 8.50 using 1 M NaOH and the solution was stirred at RT for 2 hours. The reaction was quenched by adjusting the pH to 4.0. The solution was then dialyzed (MWCO 14 kDa, regenerated cellulose,

ThermoFisher Scientific) against pH 4.00 DI-$H_2O$ for 3 days while protected from light. Dialysis water was refreshed twice daily. Purified HA was passed through 0.22 µm filters (EMD Millipore), flash frozen using liquid nitrogen and lyophilized. The dried product was vacuum sealed and stored at -20°C. Thiolation percentage was measured using $^1$H-NMR spectroscopy and an Ellman's assay for free thiols.

**Hydrogel fabrication and characterization:** Hydrogel precursor solution was prepared by dissolving HA-SH (0.5% w/v), 4-arm thiol terminated polyethylene glycol (PEG-SH) (Laysan Bio), 8-arm norbornene terminated polyethylene glycol (PEG-Norb) (Jenkem), 0.025% w/v lithium phenyl-2,4,6 trimethylbenzoylphosphinate (LAP, Sigma-Aldrich) and 0.25 mM thiolated peptides (RGD: GCGYGRGDSPG; IBSP: GCGYGGGGNGEPRGDNYRAY; JenKem, USA) in 20 mM HEPES buffer (pH=7). The hydrogel precursor solution was cast into 8 mm diameter silicone rubber molds (Grace Biolabs) and irradiated with long-wave UV (365 nm, 4.2 mW/cm$^2$) (Blak-Ray™ B-100A UV lamp, UVP™) for 15 seconds. Hydrogel storage moduli (G') were measured using a discovery hybrid rheometer-2 (DHR-2, TA Instruments) at 37 °C. Frequency sweeps were performed under 1% constant strain in the range of 0.1 to 1.0 Hz. Storage modulus of each sample was calculated as the average value of the linear region of the storage curve from the frequency sweep plot. For statistical analysis, 3 separate measurements were taken in which 5 samples from each condition were measured.

**GBM encapsulation in hydrogels**: Patient-derived GBM cell lines, HK217 (proneural), HK301 (proneural) and HK280 (mesenchymal) were used for encapsulation. Sizes of GBM spheroids were standardized by seeding approximately 600K cells per well into Aggrewell™ well plates (Stemcell Technologies) one day prior to encapsulation. The following day, spheroids were harvested from the wells, centrifuged briefly (200xG, 1 min.) and resuspended in the hydrogel precursor solution. Spheroid-laden hydrogels were formed as described in the above hydrogel fabrication section. Cell migration was observed periodically (Imaged at Day 1,3,6 and 9) by acquiring phase contrast images on a Zeiss Axio.Z1 Observer microscope with a Hamamatsu Orca Flash 4.0 V2 Digital CMOS Camera and Zeiss ZEN 2 (Blue Edition) software. Quantitation is described separately. At the end of experimental period, hydrogels were fixed using 4% paraformaldehyde (PFA) and stained with Hoescht (nuclei) and Cell Mask™ (cell membrane). These gels were imaged using a Leica SP5 confocal microscope. To block ITGαV-IBSP interaction, GBM spheroids were incubated with 10 μg/ml anti-ITGαV antibody a day prior to encapsulation. In addition, after the encapsulation, hydrogels were incubated in GBM media containing 10 μg/ml of the same antibody over the course of the experiment.

**Migration analysis in hydrogels:** Cell migration was quantified using shape factor (circularity) and length of migration from sphere edge. To calculate circularity, perimeter of each sphere was marked in ImageJ software and circularity ($4\pi A/P^2$, A=area, P=perimeter) was calculated using ImageJ shape description. Briefly, using the free drawing tool, we traced the periphery of spheres and then calculate the shape factor using shape description analysis in image J. Shape factor is defined as $4\pi A/P^2$, where A is the area and P is the perimeter of a spheroids. Shape factor values in general range from 0 to 1 in which 1 means complete circle and values smaller than 1 means deviation from a circle. With this method, more migratory spheres have shape factor values close to 0. As an arbitrary measurement, based on our data, we categorized migration in to 1) non-migratory: shape factor 1-0.7, 2) mild migration: shape factor 0.7-0.3 and 3) invasive: shape factor < 0.3. We measured shape factor for 10 spheres per group from 3 independent experiments. For average migration distance, we used the line measurement option in ImageJ and measured the length of migratory processes from the sphere periphery for 10 GBM spheroids and 3 independent experiments.

**GBM-GVC co-encapsulation:** Primary GVC line was generated from a recurrent GBM patient tumor (as described in isolation of EC by MACS). Cells were cultured in endothelial growth media (R&D systems, #CCM027) containing 2% FBS, 1% Penicillin-Streptomycin solution and growth supplement, and maintained for up to 5 passages from initial establishment. Characterization was performed by immunostaining using anti-human CD31 antibody at P1 and P3. Human brain microvascular endothelial cells (HBEC) were used as control for normal BVC. All of the ECs were used between passage 2 and 5. RNA-sequencing was performed at Passage 4 to further characterize these cultured tumor vascular cells. HBEC were cultured using standard endothelial cell media. GBM cells were infected with a lentivial vector encoding for blue fluorescent protein (BFP) to distinguish them from GVC which were infected with mCherry. We used the Aggrewell$^{TM}$ (above) to obtain spheroids of relatively uniform sizes. Co-cultures of GVC and GBM cells were established in two steps. First, GBM spheroids were resuspended in HA hydrogel precursor, casted in 4 mm diameter, silicone rubber molds and hydrogels crosslinked as described above. In the second step, GBM spheroid-laden hydrogels were transferred into 8 mm diameter molds and solution of HA-RGD (500μM RGD) containing GVC ($10^7$ cell/ml) was casted around the initial hydrogel and formed under UV. Final hydrogels were transferred to EC medium and imaged periodically (timepoints similar to previous part) as described above. At the end of experiments (Day 9), hydrogels were fixed using 4% PFA and imaged using a Leica SP5 confocal microscope.

**shRNA Screening:** shRNA clones of the genes of interest were arrayed from our genome wide shRNA library (Silva et al., 2005). 100 ng shRNA encoding pGIPZ plasmid was spotted into PDL coated 96 well plates and 100 ng pCMVd8.91 with 10 ngpMD2G was added together with Mirus TransIT in a 1:3 ratio and incubated for 20 min in a total volume of 25 µL. 75 µL containing 40,000 293T cells were added on top to a total of 100 µL in DMEM with 30% FBS with 1x PSG, NEN and HEPES. Successful transfection was confirmed after 24 hours at 37°C and 5% $CO_2$ by

GFP expression of the cells. Virus was harvested after 48 hours at 37°C and 5% $CO_2$ and 7.5 µL virus containing media was plated into each well of a 384 well plate using an Agilent Vprep in a custom HEPA filtered enclosure. Media was added to 25µL total volume before cells were added at [100000/ml], to a total of 50ul volume/well. 3 days post transduction, small GBM spheroids were encapsulated in hydrogels plated in another 384-well plate.

**High-throughput imaging and quantification of gliomasphere migration:** Gliomaspheres were encapsulated in hydrogels and plated in 384-well Greiner plates and imaged using a Molecular Devices ImageXpress XL platform. In short, plates were imaged using a Nikon 10x objective (0.3NA, Plan Apo) with no binning and laser auto-focusing. Plates were imaged daily, and were treated with Hoechst, at 1:3000 in media, overnight before the final imaging that were to be used for quantification. The resulting images were analyzed using the MetaXpress Custom Module editor. A custom module was set up using Adaptive Thresholding in the UV/DAPI channel with a size window from 10-400m micron and an intensity over local background of at least 1750 grey scales. This analysis applied a mask to the images, thus allowing for quantification of the number of objects dispersed within the hydrogel. Though this method of quantification is an underestimation of the number of cells, it does provide for an efficient means to quantify dispersion from the gliomasphere core for our HT purposes. The following parameters were extracted on an object by object base: Total area average, area average per object, centroid position for x and y axis. Also, the sums for the same parameters were extracted. The Elledge form factor was extracted on an object by object base.

**Fluorescent in situ hybridization by RNAScope:** FFPE tumor sections were baked at 60℃ for 1 hour, then deparaffinized with xylene and rehydrated in a series of washes with decreasing ethanol concentrations. Antigen retrieval and *in situ* hybridization was performed according to the protocol for the RNAScope™ Fluorescent Multiplex Assay (ACDBio). Targets were labelled with Cy3- and Cy5-tyramide TSA solutions (1:700, Perkin Elmer) and coverslips mounted with Prolong Gold Mounting Medium with DAPI.

**Cell Proliferation Assay:** The Cell Counting Kit-8 (Dojindo Molecular Technologies Inc.) was used according to the manufacturer's protocol. Freshly dissociated gliomaspheres were plated at 5000cells/100µl/well in a 97-well plate and allowed to proliferated for 7 days at which point cell numbers of the experimental conditions (250nM IBSP in all cases unless specified otherwise) were assessed as compared to control (CTRL) conditions.

**Western blot:** Protein lysates were obtained from 300k cells per sample using RIPA buffer supplemented with protease and phosphatase inhibitors. To prepare western blot samples, protein solutions were mixed with Laemmli buffer (2X, contain 5 %v/v β-mercapto ethanol, bio-rad) in 1:1 ratio and heated at 97 °C for 5 minutes. Samples were loaded in a Nupage™ 4-12% bis-Tris protein gel (Thermoscientific). Gels were run in MOPS-SDS buffer (20X, thermo fisher) at 60V for 15 minutes the 165V for 1 hour. Later proteins were transferred onto a PVDF membrane (Thermo Scientific™) in tris/glycine (10X, Bio-Rad) buffer containing 20 %v/v methanol. IBSP detection was done using a human IBSP polycolonal antibody (Rabbit, Thermo Fisher Scientific) as the primary antibody and then a goat anti-rabbit IgG (HRP linked, cell signaling). For the housekeeping gene, GAPDH was stained using a GAPDH loading control antibody (mouse, Fisher Scientific) and then a goat anti-mouse IgG secondary antibody (HRP, Novus biological). Protein bands were developed using the Clarity™ western ECL substrate (Bio-Rad). Protein bands were visualized using MYECL gel imager (Thermo Scientific). Quantification of protein levels were done in Image J.

**Ex-vivo IBSP treatment and generation of murine tumor xenografts:** 6-8 weeks old SCID mice were used in this study and experiments were carried out under an Institutional Animal

Care and Use Committee (IACUC) approved protocol according to NIH guidelines at UAB. GBM cells derived from patients were pretreated for 3 days in synthetic IBSP or control peptide (IBSP-SCR); 10ug/ml; 0.33mM. On the day of injection, concentration of IBSP was increased 3x to 30ug/ml (1µM) in injection solution. 200,000 GBM cells were injected into the brains of SCID mice. Four weeks after injection, mice were sacrificed and perfused with ice-cold PBS and 4% (wt/vol) paraformaldehyde (PFA). Brains were dissected and fixed in 4% PFA for 24, hours and then transferred to 10% formalin, and sectioned for staining.

**IBSP WT and KO mice tumor transplants and imaging:** 50,000 cells from an IDH WT

murine glioma line (Nunez et al., 2019) firefly-luciferase was transplanted into 8-12 weeks old *Ibsp* Wt and Ko mice. One week following transplantation, tumors were imaged for luciferase signal using IVIS Lumina II imager at the Crump Institute's Preclinical Imaging Technology Center at UCLA. Briefly, animals were anesthetized with isoflurane and injected intraperitoneally with D-luciferin (100µl; GoldBio) dissolved in phosphate buffer saline without Ca2+ or Mg2+ (30 mg/ml). 4 minutes after injection animals were imaged on an IVIS Lumina II (Caliper Life Sciences for 4 minutes. Bioluminescence images were overlaid on photographs of the mice using Living Image software (Perkin Elmer) and ROIs were selected to encompass the tumor area and radiance was used as a measure of tumor burden.

**Tumor Cell-Blood Vessel Distance Quantification:** 30µm thick cryosectioned tissue sections containing GFP-expressing tumors were stained with Hoechst nuclear stain (Millipore Sigma) and DyLight 649-conjugated tomato lectin dye (Vector Labs) to label blood vessels. Images were acquired on an AxioImager.M2 (Zeiss) equipped with an Apotome 2. Pseudoconfocal z-stacks were taken to envelope the depth of the tissue section and a maximum intensity projection was used for quantification. File names were de-identified and randomized for ImageJ analysis by blind observers. A border was drawn around the main tumor and only cells outside the border were counted and measured. Distance in μm between GFP-expressing tumor cells and nearest lectin-stained blood vessel were measured using the line tool. Data is represented as boxplots of individual cell-blood vessel measurements for each animal.

**Statistics:** Small group comparisons were done using the Student's t test, where significance was determined at $p<0.05$ unless stated otherwise. All error bars on the graphs shown are a measure of the Standard Error of Mean (SEM) or Standard deviation (SD) unless specified otherwise. Gene expression and correlation statistical analysis methods are described above. For gliomasphere migration in HA 3D culture systems, normality of each data set was analyzed using D'Agostino & Pearson omnibus normality test. For normally distributed population, one-way ANOVA followed by post-hoc t-test were used to determine the statistical differences among the groups. For normal distributions, Mann-Whitney non parametric test was used.

**Supplementary Files:**

**File S1:** Complete overall GTC vs. glial progenitor cell (BGPC) differential expression analysis (DEA) (a), and the associated Ingenuity Pathway Analysis (IPA) Diseases and Functions $p<0.05$ (b);

**File S2:** Complete overall GVC vs. BVC DEA (a), the significantly enriched IPA Canonical Pathways, $p<0.05$ (b), and the IPA-predicted Upstream Regulators, $p<0.05$ (c);

**File S3:** GVC extracellular factors DEA (a), and GTC plasma membrane (PM) proteins DEA (b);

**File S4:** GVC extracellular factors FPKM expression units (a), GTC PM proteins FPKM expression units (b), and our comprehensive PVN interactome by FPKM expression units (c), where FPKM>4 are highlighted in red

**File S5:** Complete overall IBSP vs. Control DEA, FDR<0.001 (a), the significantly enriched IPA Canonical Pathways, $p<0.05$ (b), the IPA-predicted Upstream Regulators, $p<0.05$ (c), and associated IPA Diseases and Functions, $p<1.5e-6$ (d).

**Supplementary figures:**

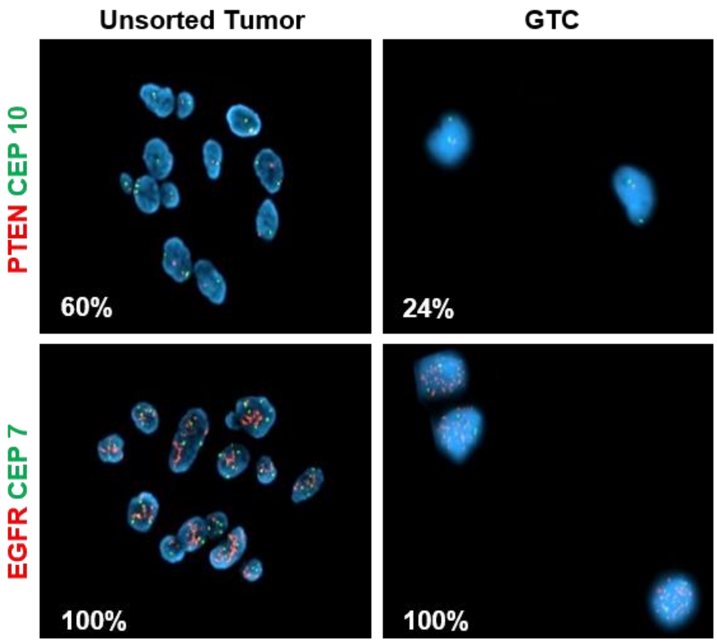


| Gene | FC GTC vs. GBM | p-Value | FDR |
|---|---|---|---|
| OLIG2 | 3.87 | 7.38E-03 | 3.39E-02 |
| CSPG4 | 8.79 | 3.13E-07 | 1.39E-05 |
| SOX10 | 8.15 | 1.25E-03 | 8.84E-03 |
| NKX2-2 | 6.10 | 8.84E-03 | 3.88E-02 |
| STAT3 | 2.53 | 1.57E-03 | 1.05E-02 |
| KLF4 | 22.59 | 1.10E-19 | 3.35E-16 |
| GFAP | 2.87 | 4.64E-03 | 2.40E-02 |
| MSI1 | 6.44 | 8.07E-04 | 6.26E-03 |
| POU5F1 | 3.93 | 2.61E-02 | 8.74E-02 |
| **Gene** | **FC GTC vs. BGPC** | **p-Value** | **FDR** |
| NES | 9.74 | 1.94E-03 | 2.67E-02 |
| SALL4 | 111.22 | 6.98E-03 | 5.84E-02 |
| POU5F1 | 49.82 | 1.81E-02 | 1.04E-01 |
| MYC | 8.20 | 4.12E-03 | 4.22E-02 |

B

| DEA GTC vs. BGPC: Diseases or Functions Annotation | z-score | p-Value |
|---|---|---|
| proliferation of cells | 4.19 | 7.3E-05 |
| cell survival | 2.91 | 2.2E-03 |
| growth of malignant tumor | 2.89 | 6.3E-04 |
| gliomatosis | 2.88 | 6.5E-04 |
| proliferation of tumor cells | 2.85 | 4.6E-03 |
| central nervous system tumor | 2.69 | 1.7E-04 |
| neuroepithelial tumor | 2.41 | 4.6E-04 |
| segregation of chromosomes | 2.36 | 6.3E-05 |
| apoptosis of vascular endothelial cells | 2.33 | 3.9E-03 |
| Congression of chromosomes | 2.24 | 4.3E-05 |
| homologous recombination | 2.17 | 4.4E-04 |
| interphase of colon cancer cell lines | -2.38 | 5.9E-04 |
| learning | -2.53 | 7.2E-04 |
| cognition | -2.65 | 3.8E-03 |
| spatial learning | -2.80 | 1.9E-03 |

D

| Expression Intensity (FPKM) | | | |
|---|---|---|---|
| **Gene** | **GBM** | **GTC** | **BGPC** |
| CSPG4 | 0.16 | 2.20 | 1.99 |
| OLIG2 | 4.33 | 27.47 | 35.56 |
| SOX10 | 3.96 | 39.38 | 61.74 |
| NKX2-2 | 0.38 | 4.55 | 4.30 |
| STAT3 | 2.19 | 6.17 | 2.99 |
| SOX2 | 14.86 | 51.17 | 20.77 |
| KLF4 | 0.59 | 17.61 | 36.56 |
| PROM1 | 0.25 | 1.29 | 0.60 |
| CD44 | 4.33 | 4.34 | 1.79 |
| GFAP | 57.04 | 182.36 | 107.95 |
| FUT4 | 1.51 | 3.08 | 1.07 |
| MSI1 | 0.40 | 2.98 | 1.65 |
| MSI2 | 4.97 | 13.59 | 7.85 |
| ITGA6 | 12.00 | 7.67 | 2.29 |
| NES | 12.73 | 9.42 | 0.89 |
| MYC | 1.72 | 3.23 | 0.30 |

**Figure S1: Characterization of the A2B5+ tumor cell population**

(A) Cytogenetic analysis of A2B5+ GTCs and their parent tumor (clinical case) is shown. Dual color FISH was performed as per the UCLA Clinical Pathology protocol for chromosome 7 and 10 centromeric regions (green) and 7P12 for EGFR or 19q13 for PTEN (red).

(B) Diseases and Functions significantly associated with GTC differential expression as compared to non-neoplastic BGPC. Only significantly activated functions as predicted according to Z-score>2 are shown ($p<0.005$).

(C) DEA of GTC vs. BGPC of various known brain cell markers

(D) intensity expression values of known brain cell markers in whole GBM samples, GTC, and BGPC.

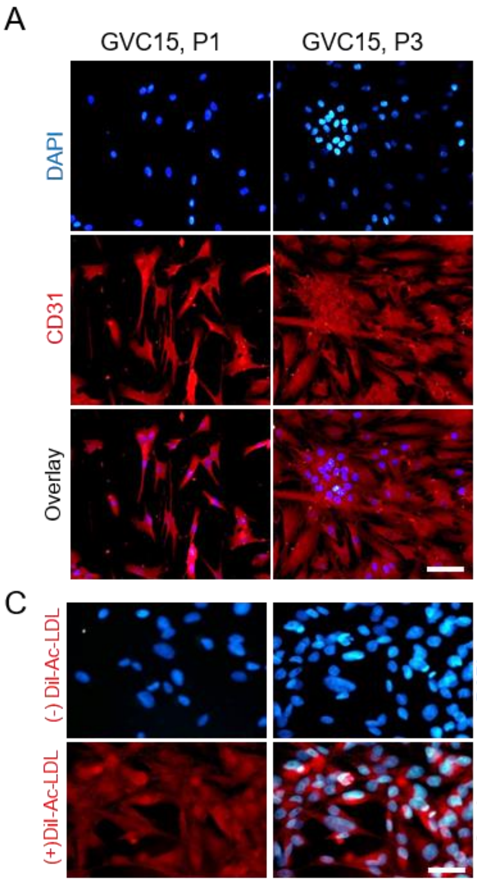


B

| | Gene | FPKM GVC15 (P4) |
|---|---|---|
| Immune cell markers | PTPRC | 0 |
| | CD14 | 0.18 |
| | ITGAM | 0.37 |
| EC markers | PECAM1 | 0.75 |
| | **CDH5** | **2.02** |
| | **KDR** | **2.01** |
| | VWF | 0.22 |
| | **EPAS1** | **37.68** |
| | TEK | 0.86 |
| | NOS3 | 0.14 |
| | **VEZF1** | **22.02** |
| Shared EC/PC markers | **CD248** | **16.46** |
| | **MCAM** | **42.05** |
| | **ACTA2** | **7.07** |
| PC markers | **PDGFRB** | **89.85** |
| | **CSPG4** | **6.76** |
| | DES | 0.32 |
| | RGS5 | 0.09 |
| | ABCC9 | 0.01 |
| | KCNJ8 | 0 |

**Figure S2: Characterization of the CD31+ GVC population**

A) Immunostaining for CD31 (red) and DAPI (nuclei) immunostaining of cultured Passage 1 and 3 (P1, P3) GVC isolated from a GBM patient (GVC15). Scale bars, 50um

B) Table shows FPKM values of transcripts of canonical immune cell, endothelial, pericyte and markers shared by both endothelial and pericytes.

C) Images of DAPI staining and DiI-Ac-LDL (red) uptake by GVC15 cells. No LDL was added to negative control. Scale bars, 50um

GTC-GVC Interactome

A

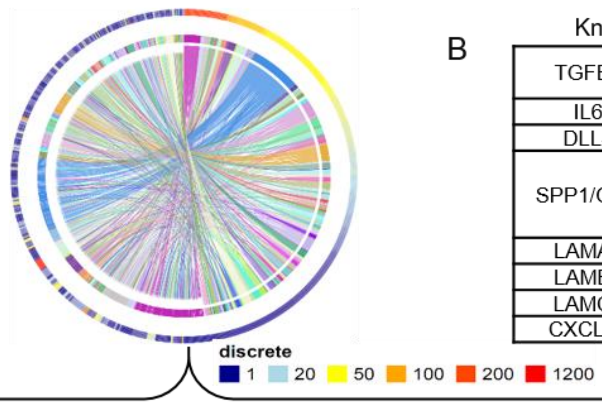


B

Known PVN angiocrine-receptor interactions

| | | | | |
|---|---|---|---|---|
| TGFB1 | TGFBR1<br>TGFBR2 | | | |
| IL6 | IL6R | | | |
| DLL4 | NOTCH1 | | | |
| SPP1/OPN | ITGAV<br>ITGB1 | ITGB3<br>ITGA5 | ITGA4<br>ITGA9 | CD44<br>EGFR |
| LAMA4 | ITGA6 | | | |
| LAMB1 | ITGA6 | | | |
| LAMC1 | ITGA6 | | | |
| CXCL12 | CXCR4 | | | |

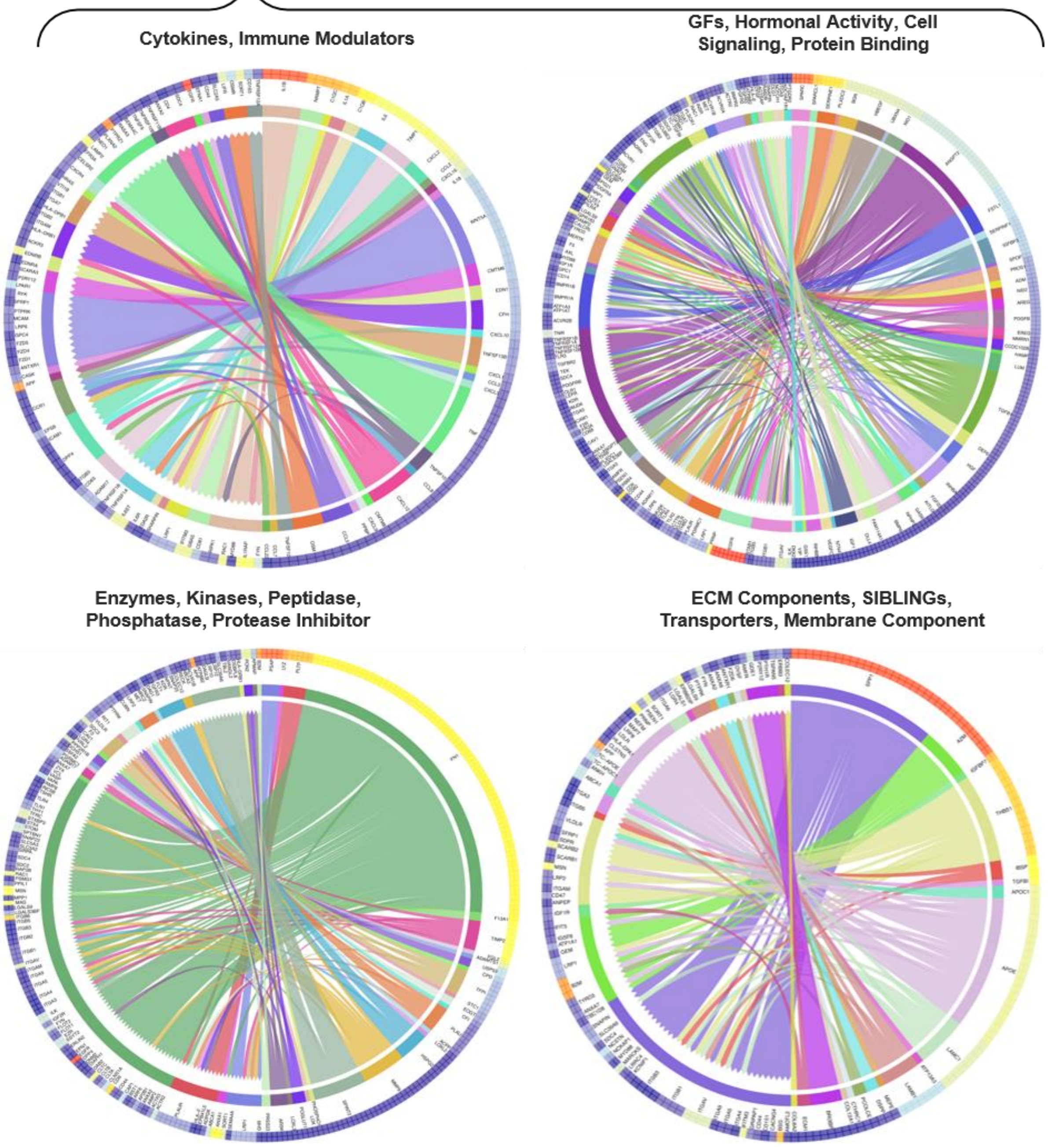

**Figure S3: GTC-GVC Interactome**

(A) Circos plots of the interactome for various processes: a) GFs, hormonal activity, cell signaling, and protein binding, b) Cytokines, Immune Modulators, c) Enzymes, Kinases, Peptidase, Phosphatase, Protease Inhibitor and d) ECM Components, SIBLINGs, Transporters, Membrane Components.

(B) Table shows known PVN ligands and their putative interactions with GTC within the interactome according to FPKM expression units

A

| Gene | Small integrin-binding ligand N-linked glycoprotein (SIBLINGs) Family of proteins | FPKM GVC | Enrichment Factor (LogFC GVC vs. BVC ) | P-Values | Candidate GTC Receptors |
|---|---|---|---|---|---|
| SPP1 | secreted phosphoprotein1 (Osteopontin) | 663.62 | 3 | 0.0008 | ITGAV ITGB3 ITGB1 ITGB5 ITGA6 ITGA9 ITGB8 ITGA1 ITGA7 ITGA5 CD44 |
| IBSP | integrin-binding sialoprotein | 50.09 | 13 | 0.00001 | |
| DMP1 | dentin matrix acidic phosphoprotein 1 | 3.22 | 8 | 0.002 | |
| DSPP | dentin sialophosphoprotein | 3.21 | 6 | 0.008 | |
| MEPE | matrix extracellular phosphoglycoprotein | 3.30 | 4 | 0.004 | |

B

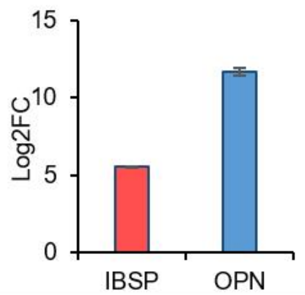


C

| Gene by Gene Correlation Significantly (P<0.001) Associated with Sphere Formation | | | | |
|---|---|---|---|---|
| Probe | Symbol | Description | Pearson correlation | P-Value |
| 212222_at | **IBSP** | integrin-binding sialoprotein | -0.72 | 1.7E-12 |
| 226513_at | **SPP1** | Secreted phosphoprotein 1 | 0.40 | 5.6E-04 |

D

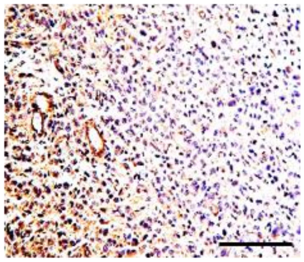


**Figure S4: IBSP is highly enriched in the GVC**

(A) Table shows the FPKM expression of SIBLING family proteins in GVC compared to BVC. IBSP is highlighted in red. SIBLING putative receptors are taken from the interactome according to FPKM expression units.

(B) Endogenous expression of IBSP as compared to OPN in primary human gliomasphere cultures (N=70).

(C) Gene expression-phenotype correlation in a gene-by-gene correlation analysis, showing IBPS and SPP1 correlation coefficients of -0.72, and 0.40 with sphere formation, respectively (p<0.001).

(D) Immunohistochemistry of IBSP (brown) in GBM patient tissue. Scale bars, 200um

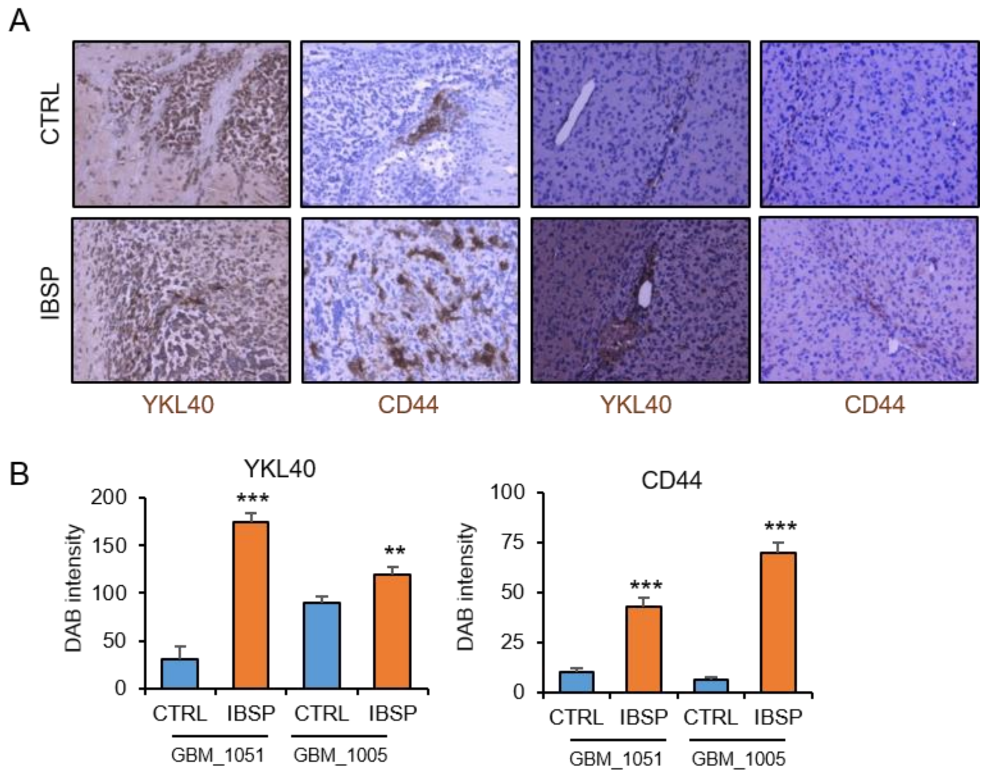


**Figure S5: IBSP promotes the expression of mesenchymal markers in tumors**

(A, B) Immunostaining of YKL40 and CD44 (brown) mesenchymal markers in CTRL and IBSP treated GBM tumors. Nuclei are counterstained with hematoxylin. Scale bars, 200um. Graphs show quantitation of average DAB intensity in each group. N=3 mice, ** and *** indicates $p<0.001$ and $p>0.0001$, t-test.

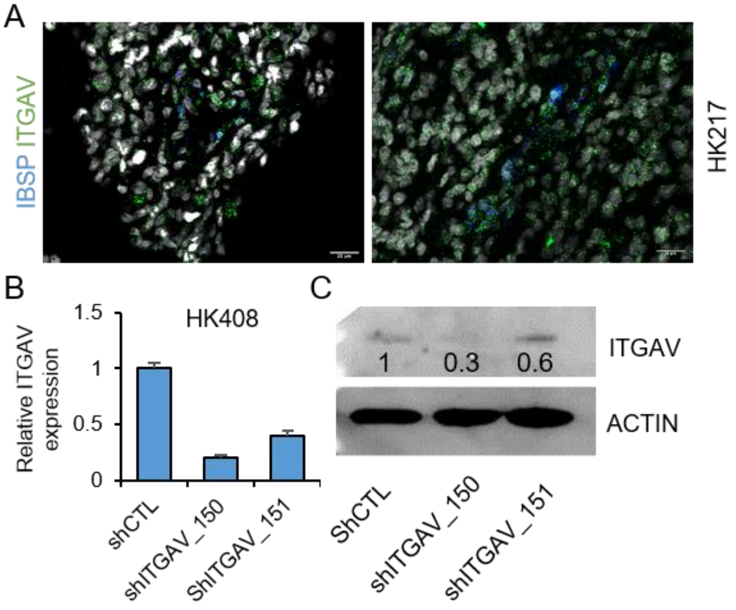


**Figure S6: ITGαV acts as the GTC receptor for IBSP**

(A) Fluorescent in situ hybridization of IBSP (blue) and ITGαV (green) in GBM tumor tissue (HK_217). Scale bars, 20um.

(B, C) ITGαV transcript and protein expression in control (shCTL) and knockdown (shITGαV_150 and 151) in HK408 tumor line. ITGAV protein levels were normalized to actin.

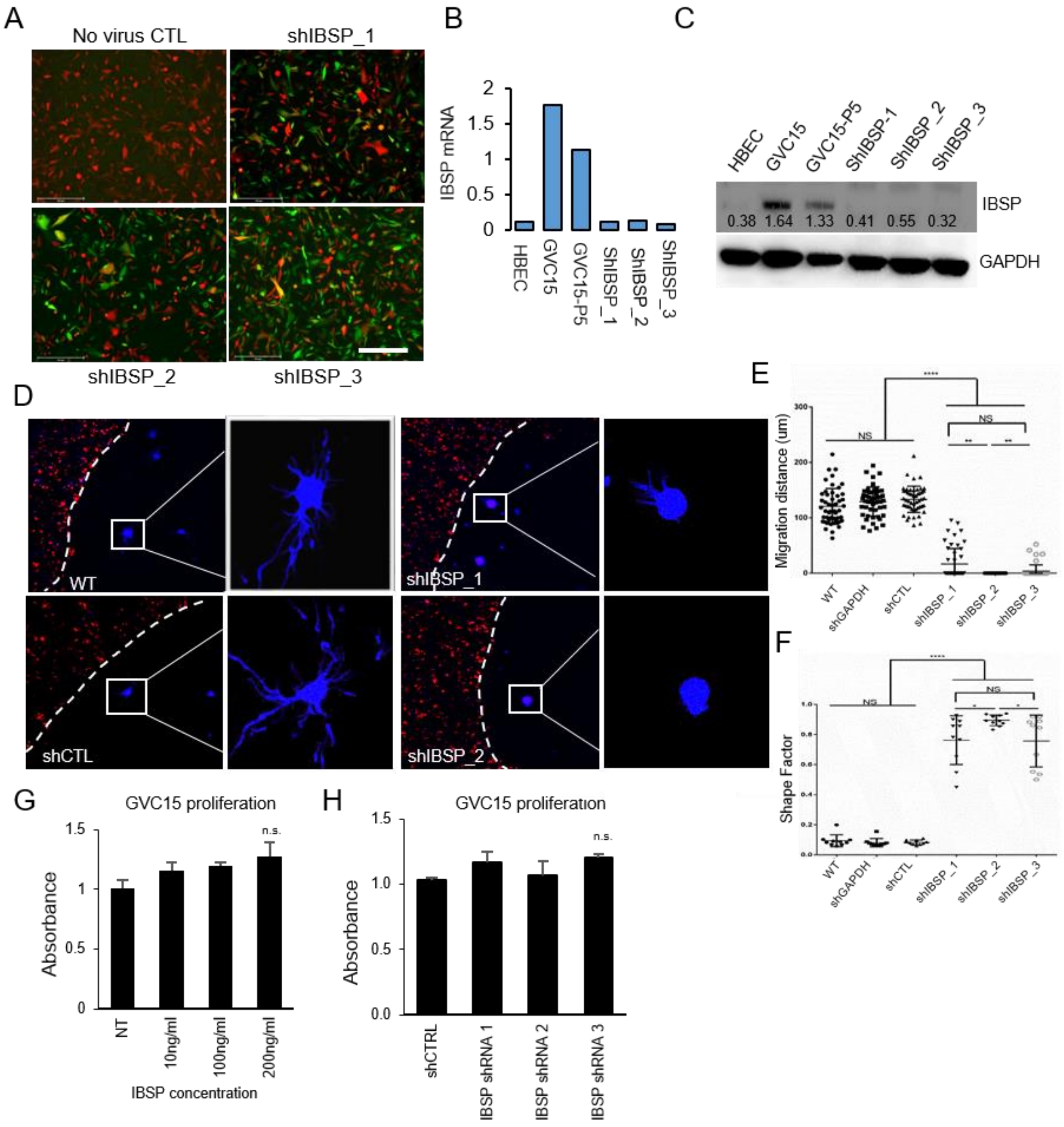


**Figure S7: Vascular-derived IBSP promotes migration of tumor cells**

(A) Images show expression of shRNA_IBSP constructs expressing GFP in GVC-infected with mCherry. Scale bars, 100um.

(B, C) IBSP transcript and protein levels in normal EC (HBEC), control GVC (P3 and P5), and IBSP-knockdown GVC15. Protein levels normalized to GAPDH is shown on the blots.

(D) Images show GVC-mCherry containing WT, shCTL or shIBSP knockdown lentivirus and GBM spheroids (blue, BFP) embedded in hydrogels. Scale bars, 250um. White dashed lines demarcate the GVC and GBM hydrogels. Boxes show the GBM spheroids at higher magnification. Scale bars,100um.

(E, F) Graphs show quantitation of migration distance and shape factor in each group. N=10 GBM spheroids, and 3 independent experiments. *** $p<0.001$, one-way ANOVA and t-test.

(G) Graph shows quantitation of GVC15 cell proliferation in response to IBSP at the indicated concentrations. Average absorbance derived from 3 replicates per group from two independent experiments. N.S. not significant, unpaired t-test.

(H) Graph shows quantitation of GVC15 cell proliferation in shCTL and shIBSP knockdown cells. Average absorbance derived from 3 replicates per group from two independent experiments.  N.S. not significant, unpaired t-test.
